# Gene-by-environment interactions are pervasive among natural genetic variants

**DOI:** 10.1101/2022.06.05.494888

**Authors:** Shi-An A. Chen, Alexander F. Kern, Roy Moh Lik Ang, Yihua Xie, Hunter B. Fraser

**Author notes:** Correspondence: Hunter B. Fraser. Equal Contribution (Shi-An A. Chen, Alexander F. Kern).

## Abstract

Gene-by-Environment (GxE) interactions are fundamental to understanding fitness landscapes and evolution, but have been difficult to identify at the single-nucleotide level, precluding understanding of their prevalence and molecular mechanisms. Most examples involving natural genetic variants exist at the level of entire genomes, e.g. measurement of microbial strain growth across environments, or loci encompassing many variants identified by quantitative trait loci mapping. Here, we introduce CRISPEY-BAR, a high-throughput precision-editing strategy, and use it to map base-pair resolution GxE interactions impacting yeast growth under stress conditions. First, we used CRISPEY-BAR to uncover 338 variants with fitness effects within QTLs previously mapped in different environments. We then measured 1432 ergosterol pathway variants from diverse lineages across six environments, identifying 205 natural variants affecting fitness measured in all six conditions, of which 93.7% showed GxE interactions. Finally, we examine pleiotropic cis-regulatory variants suggesting molecular mechanisms of GxE interaction. In sum, our results suggest an extremely complex, context-dependent fitness landscape characterized by pervasive GxE interactions, while also demonstrating high-throughput genome editing as an effective means for investigating this complexity.

## Introduction

An important issue in understanding complex traits is the phenomenon of gene-by- environment (GxE) interactions, wherein a genetic variant’s effect is dependent on the environment an organism is exposed to (Grishkevich and Yanai, 2013). For example, humans heterozygous for the sickle cell allele of beta-globin have a fitness advantage in environments that include malaria, and those with a lactase persistence allele have a fitness advantage when consuming dairy products (Luzzatto, 2012; Tishkoff et al., 2007). Identifying the genetic basis of such interactions is a key challenge in biology and is essential to the fields of medicine, genetics, synthetic biology, and evolutionary biology (Cardinale and Arkin, 2012; Li et al., 2019; Via and Lande, 1985).

Studies for identifying GxE generally come in two main varieties: forward and reverse genetic approaches. Forward genetic approaches leverage the association of natural variation to observed traits, which can be as simple as measuring the environmental response of different strains or species. With enough samples across multiple environments, genome-wide association studies (GWAS) can detect signals of GxE (Peter et al., 2018). Alternatively, quantitative trait locus (QTL) mapping uses genetic crosses between strains to create diverse progeny through recombination to calculate statistical signals that associate with environmental response (Bloom et al., 2013, 2019; Ehrenreich et al., 2012; Smith and Kruglyak, 2008). However, it is generally impossible to identify the specific variants underlying a GWAS or QTL peak without laborious follow- up experiments, due to insufficient mapping resolution, though crosses with tens of thousands of recombinant genotypes can resolve some QTLs to single nucleotides (Nguyen Ba et al., 2022; Rockman, 2012; She and Jarosz, 2018).

On the other hand, reverse genetic approaches such as constructing knockout libraries and measuring their effects on growth have single-gene resolution, and have been invaluable sources of information about the functions of genes in various organisms and their genetic interactions. However, most reverse genetics approaches to identify GxE interactions assay artificial alleles, such as gene knockouts or over-expression cassettes (Hillenmeyer et al., 2008; Jones et al., 2008). These generally do not reflect naturally occurring variants that contribute to phenotypic variation, so it is unknown whether GxE interactions of these alleles are relevant for understanding evolution. In some cases, reciprocal hemizygosity assays have been able to replace whole genes for dissecting QTL traits, but have not been able to separate the many variants within each gene (Smith and Kruglyak, 2008; Steinmetz et al., 2002). By using either forward or reverse genetic approaches alone, it is still a challenge to find the precise variants that underlie GxE (Ehrenreich et al., 2012).

Here, we combine the merits of forward and reverse genetics—integrating natural variation with massively parallel reverse genetic screens—to uncover variants harboring GxE interactions at the single nucleotide level. Previously, we showed that Cas9 Retron precISe Parallel Editing via homologY (CRISPEY) can achieve high efficiency precise editing, by utilizing a bacterial retron reverse transcriptase (RT) to generate multi-copy, single-stranded DNA (msDNA) from RNA templates *in nucleo* to facilitate homology-directed repair after Cas9-mediated genomic DNA cleavage (Sharon et al., 2018). To this end we created CRISPEY-BAR, a platform for creating and monitoring thousands of genetic variants in a single experiment. This is achieved through multiplexed, programmed installation of a predefined variant and an associated non-random barcode using a dual-CRISPEY design. Importantly, this design has improved statistical power to detect fitness effects by incorporating unique molecular identifiers (UMIs), as well as the ability to maintain strain barcodes in non-selective media, enabling us to both assay and detect GxE effects of thousands of individual genetic variants in any growth condition.

This approach allows us to survey natural variants throughout the genome in any condition, giving us the ability to decipher the precise genetic basis and molecular mechanisms giving rise to complex traits.

We used CRISPEY-BAR to measure the effects of 4184 natural variants segregating in yeast (*Saccharomyces cerevisiae*) across a variety of conditions. We pinpointed 548 variants underlying variation in growth in these environments. Importantly, the resolution of our measurements can differentiate the effects of variants even when they are tightly clustered in the genome, as well as different alleles at the same genomic position. This single-nucleotide resolution of GxE interactions not only allows us to explore the natural landscape of complex traits, but also provides direct mechanistic insights into phenotypic evolution (Lee et al., 2014; Rockman, 2012). More generally, we have established a paradigm for studying genetic variants and their environmental interactions at unprecedented resolution and throughput via multiplexed precision genome editing.

## Results

### CRISPEY-BAR enables high-resolution mapping of genotype to phenotype relationships

CRISPEY-BAR is a scalable system for measuring the effects of precise genome edits by tracking an associated genomic barcode (**Figure 1a**). As described in our previous report, CRISPEY uses a single guide/donor pair to make one precise edit per cell, and in a pooled assay, measures the change in abundance of each guide/donor pair post-editing through high-throughput sequencing of plasmids (**Figure 1b**) (Sharon et al., 2018). We developed a new vector design incorporating two consecutive retron-guide cassettes flanked by three self-cleaving ribozymes, allowing simultaneous generation of two guide/donor pairs for making two precise edits in the same cell (Riccitelli et al., 2014) (**Figure 1a, Supplemental Figure 1**). The different ribozymes prevent unwanted recombination events during pooled cloning and co-transcriptionally separate the two retron-guide RNAs for processing by retron reverse transcriptase (RT). CRISPEY-BAR implements a dual-edit design to simultaneously 1) integrate a unique genomic barcode and 2) make a precise variant edit of interest. Each variant editing guide/donor pair is associated with a unique barcode, which can be used to track change in the abundance of cells edited by a specific guide/donor pair (**Figure 1c**). Importantly, we further linked UMIs to each barcode, to serve as biological replicates for pooled-editing and growth competition (**Figure 1c**). We designed CRISPEY-BAR to measure the fitness effect of each variant with at least two guide/donor pairs, six UMIs and three pooled competition replicates (**Figure 1c, Supplemental Figure S2**).

**Figure 1:**
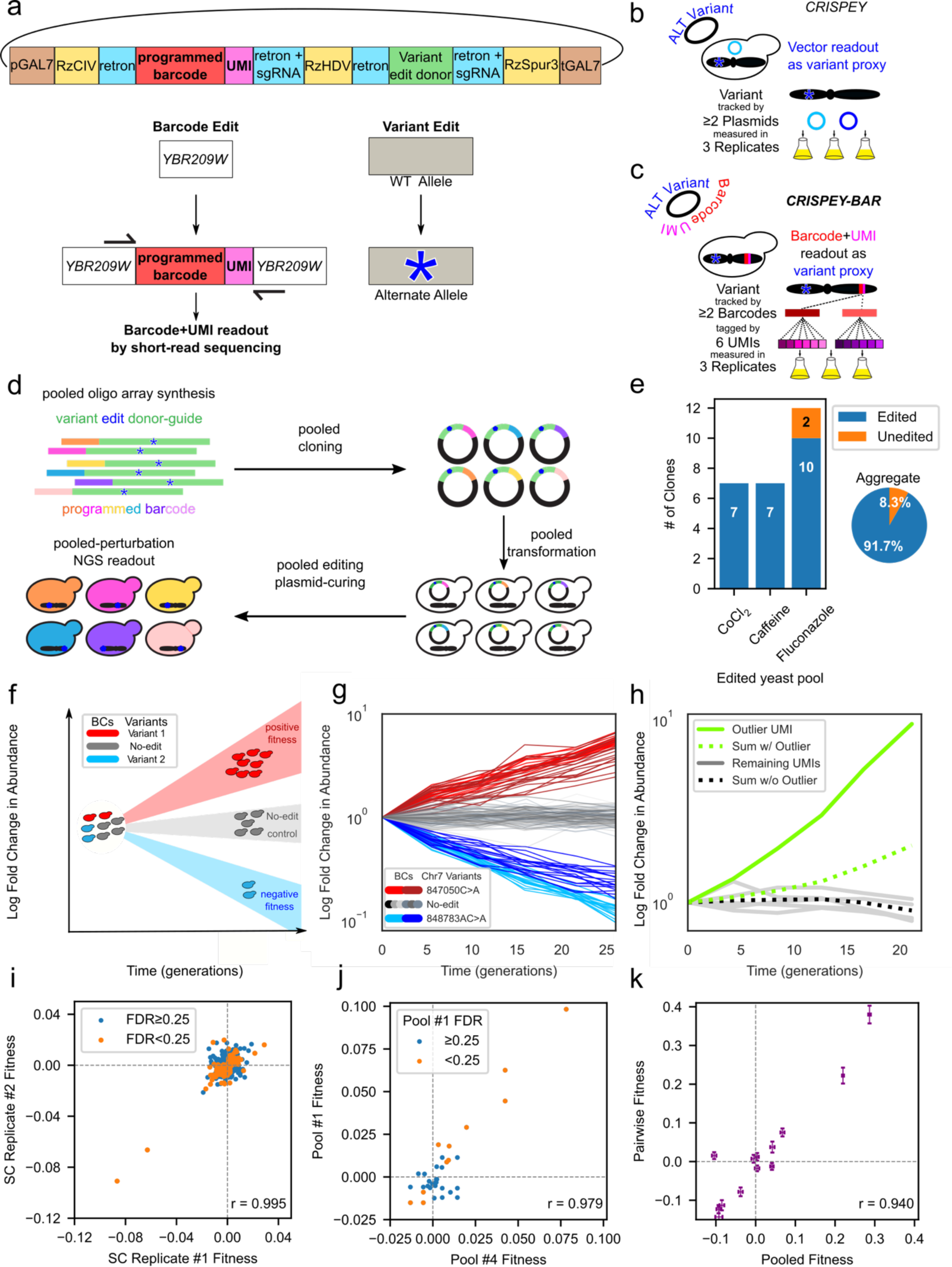
Design and validation of CRISPEY-BAR for generating and tracking thousands of precise genome edits simultaneously. *a,* Schematic of CRISPEY-BAR dual edit strategy. Top, CRISPEY-BAR expression cassette consisting of pGAL7 galactose-inducible promoter and terminator (brown); self-cleaving HDV-like-ribozymes RzCIV, RZHDV and RZSpur3 (magenta); barcode insertion retron-guide cassette (blue) containing programmed barcode (orange) and UMI (yellow); variant editing cassette (green). Middle, the variant editing cassette converts a wildtype (WT) allele into an alternative allele. Bottom, the barcode insertion retron-guide cassette *b,* Schematic for conventional CRISPEY. Variants tracked across three growth replicates by plasmids containing guide-donor oligo. *c,* Schematic for CRISPEY-BAR. Variants tracked across three growth replicates by genomically-integrated barcodes with attached UMIs. *d,* Workflow for CRISPEY-BAR library pool construction. *e,* Validation of genomic variant editing rate from CRISPEY-BAR. Blue, randomly picked colonies that contain both genomic-integrated barcode and the designed edit. Orange, randomly picked colonies that contain only the genomic- integrated barcode but not the designed edit. *f,* Schematic for CRISPEY-BAR pooled competition in yeast. *g,* Example of CRISPEY-BAR data over time. Each line indicates normalized counts for a single UMI for a given barcode from 1 of 3 replicates in a competition experiment. Counts in later time points are normalized to the first time point. Light blue and blue: two barcodes representing different guides targeting the same variant chr7: 848783 AC>A. Red and dark red: two barcodes representing different guides targeting the same variant chr7: 847050 C>A. Gray scale: Non-targeting of variants, barcode integration only (no-edit control regarding variants). Data shown are from Terbinafine competition across approximately 26 generations. *h,* Example of outlier removal. Green solid line, normalized reads from an outlier UMI. Green dotted line, normalized sum of reads from all UMIs of the barcode. Gray solid line, normalized reads from non-outlier UMIs. Black dotted line, normalized sum of reads from all UMIs of the barcode excluding outlier UMI. *i,* Replication of fitness effects between two competition triplicates in synthetic complete media (SC). Orange, variants with FDR < 0.25 in both triplicates. Blue, variants with FDR >= 0.25 in one or more triplicates. *j,* Replication of fitness effects. X-axis and Y-axis indicate fitness effects measured by two independent CRISPEY-BAR experiments, pool#1 and pool#4, in cobalt chloride. Or- ange, variants with FDR < 0.25 in pool#1. Blue, variants with FDR >= 0.25 in pool#1. *k,* Validation of pooled fitness in fluconazole by pairwise competition. X-axis, fitness ef- fect measured by CRISPEY-BAR pooled competition. Y-axis, fitness effect measured through pairwise competition against GFP strain using flow cytometry. Data shown for 13 variants in fluconazole. Data presented as mean ± SEM.

Since the barcode is genomically-integrated, no maintenance of an ectopic vector is needed post-editing, and 1:1 stoichiometric measurement of edited strains can be achieved through multiplexed sequencing of barcode amplicons (**Figure 1d**). In particular, the barcode was designed to be covered by 76-base short-read sequencing to minimize sequencing costs and run-time, instead of resequencing the plasmid with 300- base paired-end reads to re-identify guide-donor pairs (**Supplemental Figure S3**). This sequencing design uses primers that are specific to the barcode-integrated genomic locus, therefore sequencing only the barcoded strains (**Supplemental Figure S3**). Selective detection of the integrated barcode edit guarantees the edited cell expresses functional Cas9 and retron components, as well as endogenous cellular factors that facilitate HDR. This strategy allowed us to enrich for strains likely containing variant edits, which is crucial for high-throughput screens. A similar co-CRISPR strategy has been shown to improve edited mutant selection by co-injection of multiple editing vectors for both non-selectable and selectable-markers (Kim et al., 2014). We found an aggregate 92% pooled editing rate from randomly picked barcoded strains (**Figure 1e**).

The genome-integrated barcodes from a multiplexed CRISPEY-BAR library enabled us to track the abundance of thousands of programmed mutants in non-selective media (**Figure 1f,g**). Importantly, we included no-edit controls that do not install any variants apart from barcode integration to establish neutral fitness levels that arise from experimental noise and genetic drift (**Figure 1f,g**). Six pre-defined unique molecular identifiers (UMIs) were incorporated for every barcode-variant edit combination to increase biological replication, allowing us to capture the noise from variable editing rates due to guide efficiency as well as outlier detection of random mutants arising during transformation, editing, and competition to improve our estimates of variant fitness effects (**Figure 1h**). Spontaneous mutations with strong positive fitness effects in particular would be expected to dominate the reads for a given UMI, so by removing these potential outlier UMIs, we sought to reduce false positives (**STAR Methods**). We reasoned that it is unlikely that random mutations would arise for all UMI replicates for a given variant edit independent of CRISPEY-BAR.

We were able to show that CRISPEY-BAR measured fitness effects are highly reproducible between growth competition replicates. Variant fitness is approximated by fitting a linear model for estimating log2 fold-change abundance of each barcode-UMI over generation time during growth competition, as described previously (Pearson r = 0.9996, p = 1.38x10^-16^ for variants with FDR<0.25 in both replicates) (**Figure 1i,** see also **STAR Methods**) (Sharon et al., 2018). Importantly, across four independently generated and measured variant pools, measured in 13 competitions, only four putatively non-targeting barcodes had significant fitness effects in any competition at FDR<0.01, out of 43 in each pool, showing that CRISPEY-BAR has a low false positive rate. In addition, CRISPEY-BAR measured fitness effects are highly reproducible between experiments. Overlapping variants from two separately cloned, transformed, edit-induced, growth- completed, library-prepared, and sequenced CRISPEY-BAR experiments showed high replication (Pearson r=0.979, p =1.52x10^-7^ for all overlapping variants) in fitness effects for growth in cobalt chloride, despite being competed against an otherwise separate library other than technical controls (**Figure 1j**). This result shows that with overlapping variants between CRISPEY-BAR libraries, we can potentially scale pooled screening strategies with minimal batch effects. Finally, we validated 13 genotyped strains edited by CRISPEY-BAR and performed pairwise competitions in fluconazole versus a fluorescently labeled un-edited strain. The variant fitness measured by these pairwise competitions showed a high correlation with fitness measured in pooled competitions (Pearson r = 0.940, p = 1.80x10^-6^) (**Figure 1k**). In sum, CRISPEY-BAR is highly efficient in precision editing and allows massively parallel tracking of variant fitness effects using the dual-edit design.

### Detection of natural variants affecting fitness within QTLs reveals hidden genetic complexity

To evaluate CRISPEY-BAR as a high-throughput, scalable platform to measure variants’ effects on phenotypes, we first sought to characterize variants within regions likely to be enriched for effects on growth in response to stress conditions, in which the yeast pool has slower growth overall. We targeted a total of 36 genomic regions overlapping QTLs for growth of segregants derived from 16 diverse parental strains, measured in three stress conditions: fluconazole (FLC), cobalt chloride (CoCl2) and caffeine (CAFF) (**Figure 2a**) (Bloom et al., 2019). For each stress condition, we constructed a CRISPEY-BAR library pool that targets natural variants that fall within previously identified genomic regions identified by QTL mapping to affect growth in the corresponding stress condition(Bloom et al., 2019; Peter et al., 2018). We selected QTLs with 1.5-LOD confidence intervals containing only a single gene to not only increase the probability of finding fitness variants affecting fitness, but also maximize the number of QTLs surveyed given a set library size (Bloom et al., 2019). We reasoned that by installing diverse natural variants— including many not present in the 16 parental strains—we would enrich for variants impacting fitness in these stress conditions (**Figure 2a**) (Peter et al., 2018). We designed 3 oligonucleotide pools (corresponding to variants to be assayed in fluconazole, cobalt chloride, and caffeine) for pooled cloning into 3 separate CRISPEY-BAR libraries, which were then used for pooled editing (**STAR Methods**). After plasmid removal, we subjected the edited yeast to pooled growth competitions in synthetic complete media as well as each corresponding stress condition and tracked changes in barcode abundance across roughly 25 generations (**Figure 2b, Supplemental Figure S2**). To ensure the stress conditions were applied during yeast growth, we calibrated the dose of stress agents (fluconazole, cobalt chloride, and caffeine) so that the average growth rate is lower by 50% (**STAR Methods**). Importantly, we included barcodes with a non-targeting guide (designed to target a sequence which is not present in these strains) as no-edit controls to define the neutral fitness distribution within each pooled competition experiment (**Figure 1f,g**; **STAR Methods**).

**Figure 2:**
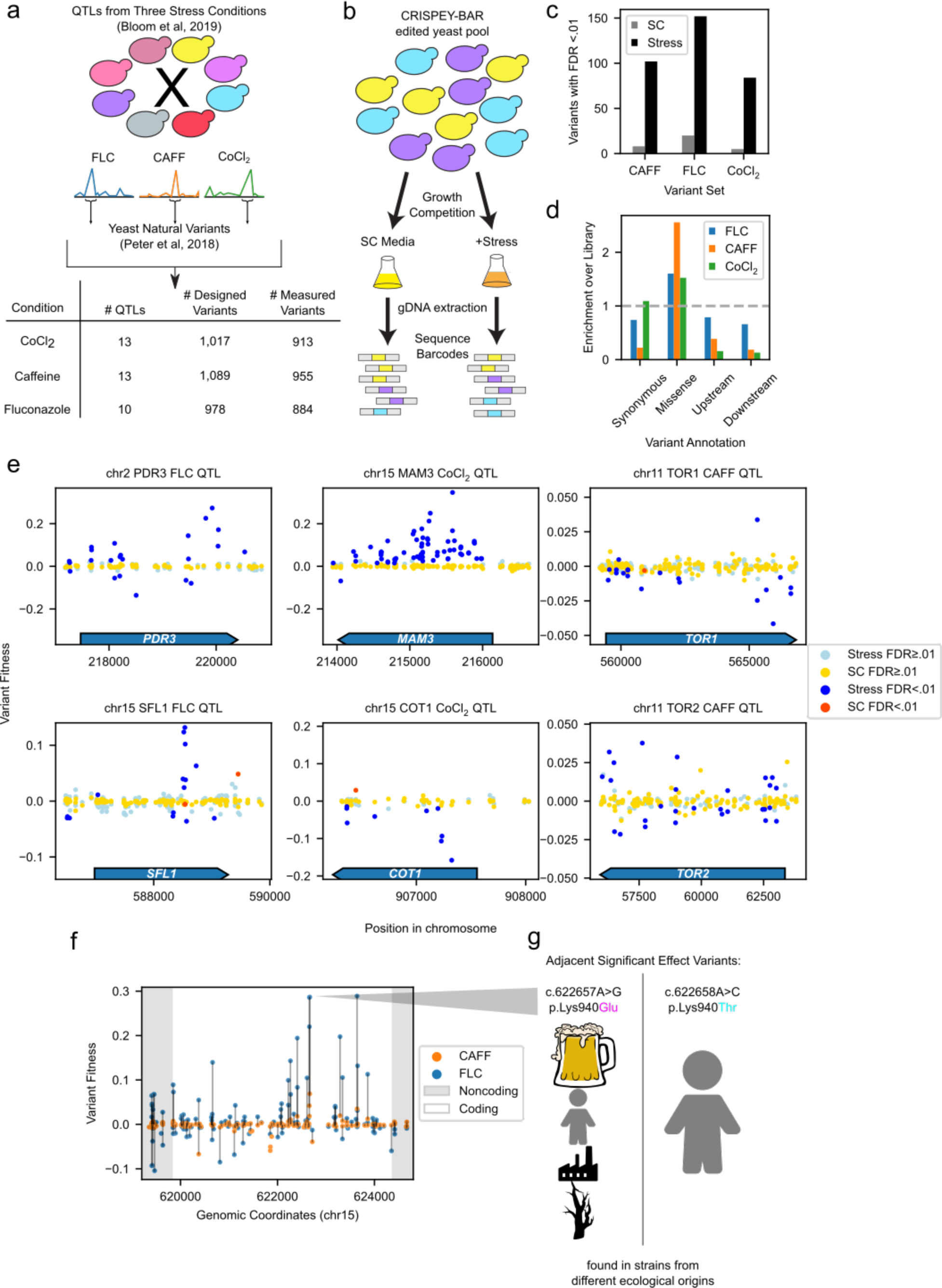
Detection of natural variants affecting fitness within QTLs mapped in complex traits. a, Diagram of library design process showing the selection of QTLs from three different conditions, sourcing of natural variants, as well as library statistics. b, Schematic for experiment workflow for QTL region variant fitness mapping with CRISPEY-BAR. c, Number of variants with fitness effect (FDR<0.01) within SC and appropriate stress condition. d, Annotation enrichment of variants with fitness effect (FDR<0.01). Blue, variant enrichment for hits in fluconazole condition. Orange, variant enrichment for hits in caffeine condition. Green, variant enrichment for hits in cobalt chloride condition. e, Fitness effects of example QTL regions. Dark blue, fitness effects in stress condition (FDR < 0.01). Dark orange, fitness effects in SC (FDR < 0.01). Light blue, no fitness effects stress condition. Gold, no fitness effects in SC. Most variants are represented twice (effect in QTL condition and complete media). f, *PDR5* fitness effects in CAFF and FLC. Magenta, *PDR5* variant fitness measured in caffeine condition. Orange, *PDR5* variant fitness measured in fluconazole condition. Dark gray, noncoding regions flanking *PDR5*. Light gray, coding region of *PDR5*. Vertical lines connect the same variant fitness values measured in both caffeine and fluconazole. g, Diagram depicting primary ecological origins of two adjacent variants mutating K940 in *PDR5* with significant fitness effects.

We identified 152 variants with significant fitness effects in fluconazole, 84 variants for cobalt chloride and 102 variants for caffeine within the regions screened for each stress condition (FDR<0.01). We found substantially fewer variants with significant fitness effects for growth in synthetic complete media (SC) than in the stress condition for each of these pools (**Figure 2c**). We next sought to identify what types of variants were most likely to have significant effects on fitness within these pools, leveraging the single-nucleotide resolution of the measurements, and saw a substantial enrichment for missense variants among causal variants in the drug conditions for all three libraries (Fisher’s exact test p= 4.72x10^-3^ for cobalt chloride, 1.20x10^-6^ for fluconazole, 1.31x10^-29^ for caffeine) (**Figure 2d**). Within many of the QTLs we identified dozens of causal variants (**Figure 2e**). For instance, 65 out of the 66 causal variants in *MAM3* for growth in cobalt chloride increased fitness, indicating that they may impair function of this gene, as *MAM3* knockout increases resistance to cobalt chloride (Yang et al., 2005). For other QTL genes such as *TOR2*, the knockout of which has been shown to decrease fitness in the presence of caffeine, we identified many variants both increasing and decreasing fitness (Reinke et al., 2006). The base-pair level resolution of CRISPEY-BAR enables us to identify substantial genetic complexity hidden within these QTLs.

One QTL gene, *PDR5*, was shared between our caffeine and fluconazole pools. This well-studied multi-drug transporter had multiple variants affecting fitness in both conditions, many of which had effects in the same direction between the conditions (**Figure 2f)** (Balzi et al., 1994; Harris et al., 2021). For example, we were able to identify two variants with fitness effect located one base-pair apart in the genome, both of which cause missense changes to the same lysine residue in *PDR5* (**Figure 2g**). These two variants both had substantial positive fitness effects for growth in fluconazole and caffeine, and do not co-occur in strains within the 1011 yeast genomes collection, indicating that they arose independently (Peter et al., 2018). One of these variants is found almost exclusively in strains with the origin “Human, clinical,” while the other is more broadly distributed across ecological origins. Beyond *PDR5*, there were several other cases where two missense variants changed the same amino acid, both having strong fitness effects (V136, G1967, and A2403 in *TOR1,* K768 in *TOR2*, G398 in COT1, and N738 in *SWH1)*.

To identify whether alleles with positive fitness effects in QTL conditions were enriched in yeast strains from particular ecological origins, we counted the number of positive effect alleles each strain in the 1,011 yeast genome strains had, including reference alleles which were beneficial relative to the negative effect engineered variants (**STAR Methods)**. Interestingly, 41 of the top 100 scoring strains are from the ecological origins “Human” and “Human, Clinical,” which is a substantial enrichment (hypergeometric p =1.41x10-13). Notably these 41 strains came from four different clades. While this result could be partially driven by population structure, it is also potentially suggestive of selection for increased fluconazole resistance among yeast isolated from human origins, warranting further study. This could be done by sampling variants from more regions of the genome, focusing on variants with high allele frequencies in particular clades for comparison.

### The GxE landscape of ergosterol synthesis pathway

Having established the ability to identify multiple natural variants affecting fitness across various environments with CRISPEY-BAR, we next examined GxE interactions within the ergosterol biosynthesis pathway (Rodrigues, 2018). This essential metabolic pathway is of great biomedical importance, being the target of multiple classes of antifungal drugs, as well as statins and has also been shown to be affected by various other stress conditions, owing to its complex transcriptional and post-transcriptional regulation (**Figure 3a**) (Bhattacharya et al., 2018; Kern et al., 2021). We sought to test natural variants within genes in this pathway as well as 1000 bp upstream and 500 bp downstream of each ORF to capture promoters and downstream regulatory regions in five stress conditions as well as SC media (**Figure 3b**). Across these six environments, we were able to capture a total of 1432 variants passing minimum read filters and outlier detection for at least one of the six conditions (**STAR Methods**).

**Figure 3:**
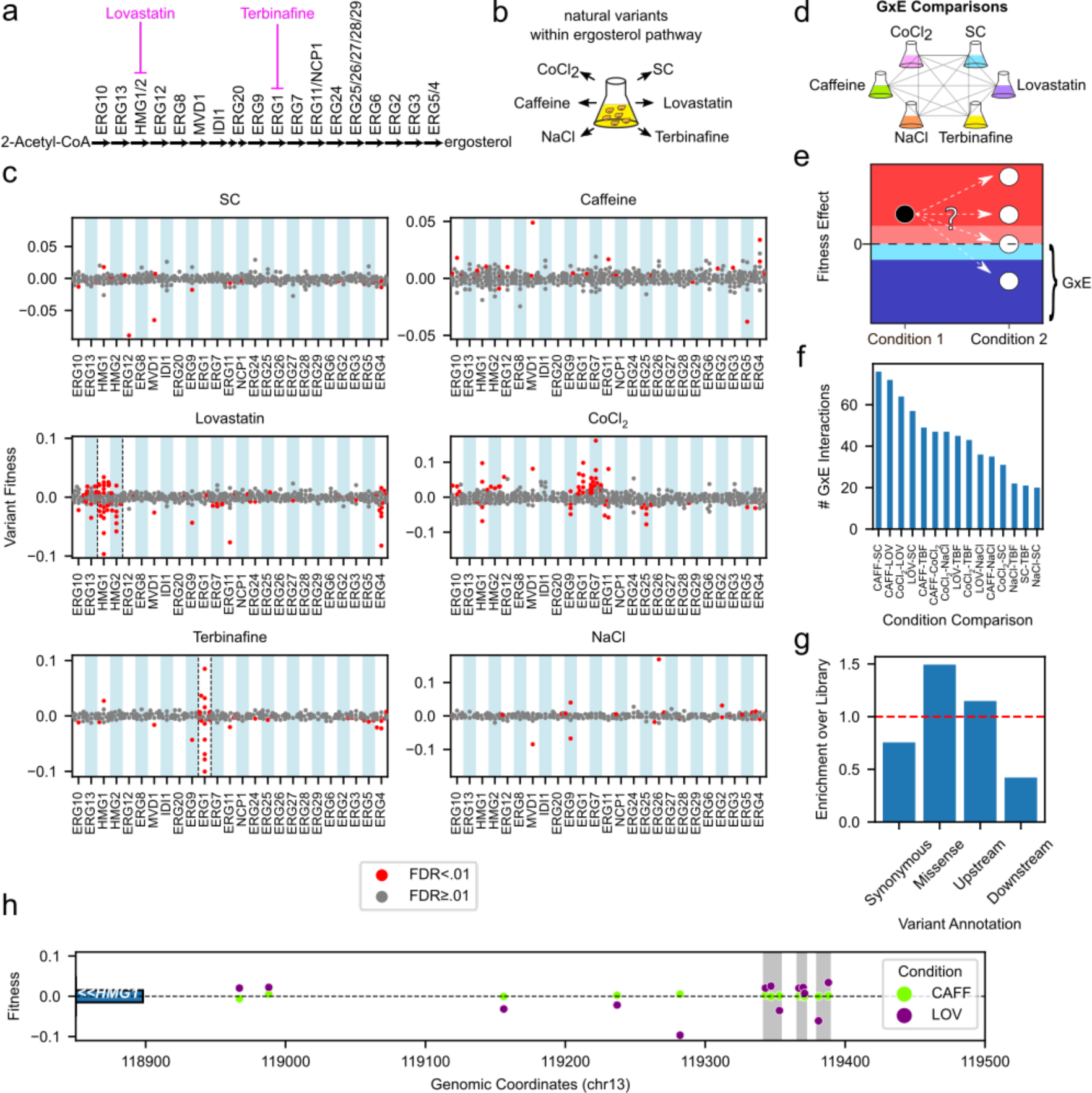
CRISPEY-BAR enabled robust mapping of variant-level GxE interactions within the ergosterol biosynthesis pathway. *a,* Ergosterol pathway diagram showing 25 genes from the ergosterol synthesis pathway surveyed in this study. Lovastatin and terbinafine target genes in the ergosterol pathway, indicated by magenta inhibition lines. *b,* The same pool of yeast edited at natural ergosterol pathway variants was grown in six different conditions and tracked by barcode sequencing. *c,* Gene level fitness effects of surveyed natural variants in six conditions. X-axis labels indicate the genes containing the variants. Red, variants with fitness effects (FDR < 0.01). Gray, non-significant variants. Target genes are outlined by dashed black lines where applicable for a given condition. *d,* GxE interactions were calculated between each pair of conditions (15 pairwise comparisons). *e,* Diagram showing definition of GxE variants in this study: A positive effect variant (black circle) in condition 1 can either have the same effect in another condition (white circle at same height in red region), a stronger positive effect (top white circle in red region), no effect, white circle at zero, or a negative effect (bottom white circle in blue region). If the variant has a negative effect or no effect at all in condition 2 (blue and light blue regions), it is labeled as GxE. *f,* The number of significant GxE interactions for each pairwise comparison. *g,* GxE annotation enrichments for variants with GxE. Enrichment of variants with GxE in each category were normalized to all variants tested. Red dashed line indicates an enrichment factor of 1.0, corresponding to no enrichment over the library. *h,* Variants with GxE effects within the *HMG1* promoter. Clusters of variants with significant GxE effects within 8 bp of each other are in gray highlighted areas. Beginning of the *HMG1* gene body is shown as a blue rectangle. Green, variants fitness effect in caffeine (CAFF) condition. Purple, variant fitness effect in lovastatin (LOV) condition.

Mapping of variants affecting fitness in our two drug conditions, lovastatin and terbinafine, revealed that the target genes for these drugs (*HMG1/2* and *ERG1* respectively) were enriched for variants showing strong fitness effects in these conditions (lovastatin p = 6.62x10^-12^ and terbinafine p =1.55x10^-16^, hypergeometric test) (**Figure 3c**) (Jandrositz et al., 1991; Lum et al., 2004; Rine et al., 1983). This illustrates the specificity of the variant effect measurements obtained from these screens. Notably, though these conditions were enriched for variants with strong effects in the target genes, variants in other ergosterol pathway genes also affected fitness, revealing extensive genetic complexity. In addition, the variants with the strongest fitness effects in the different conditions had different variant annotation enrichments. For instance, the 100 variants with the strongest fitness effects in SC were enriched for upstream/promoter variants (hypergeometric p = 3.61x10^-3^), while the 100 strongest effect variants in sodium chloride were not (hypergeometric p = 0.137).

To identify GxE variants, we performed all pairwise comparisons between the relative fitness measurements for each variant in each condition to see if the effects on growth were significantly different (**Figure 3d,e)**. To determine a reasonable threshold to define GxE variants and see if our approach to identifying GxE was robust, we performed two identical competitions in SC media and tested variants for GxE interactions between them. With the significance threshold we used (FDR <0.01 and a change in sign of fitness effect) there were zero variants showing significant GxE between the replicates, compared to a mean of 44.3 for comparisons between conditions, suggesting that our approach to identifying GxE has a low false positive rate. Since there were no significant differences between SC replicates, we combined these replicate competitions in our final analysis to increase statistical power. Overall, at this threshold we identified 256 distinct variants with at least one significant GxE interaction (GxE variants), harboring 665 pairwise GxE interactions. We next examined annotation enrichments for GxE variants and found that missense variants were strongly enriched (two-sided Bonferroni-corrected hypergeometric p = 2.09x10^-6^), while synonymous variants were depleted (two-sided Bonferroni-corrected hypergeometric p = 1.70x10^-3^) (**Figure 3f**).

The stringent definition of GxE above explicitly excludes variants with significantly significant fitness effects which are in the same direction, which we term “magnitude GxE” (red and pink in **Figure 3e**). While magnitude GxE is interesting, it also may be highly dependent on many experimental variables such as drug concentration. Thus, we opted for a stricter definition of a GxE variant: any variant with a significant GxE interaction which has measured fitness effects in opposite directions in the two conditions (blue in **Figure 3e**).

With CRISPEY-BAR, we were able to measure more than one variant at the same genomic locus for multiallelic loci within the ergosterol pathway, which highlights the resolution and specificity of the measurements. There were five multiallelic sites which had a missense variant with a significant fitness effect in one or more of the growth conditions. For three of these sites, the other variant was a synonymous variant with no effects on fitness. For the other two sites, there was one site within *HMG2* where there were two missense variants (C788F and C788Y), which had similar effects on fitness. Strikingly, the other site in *HMG1* had two missense variants making P1033A and P1033T changes which had significant effects in opposite directions on growth in lovastatin, perhaps due to the different chemical properties of threonine (polar) and alanine (nonpolar).

Next, we looked at the genomic locations of the GxE variants relative to one another to see if there was any spatial structure to GxE. We defined significant GxE variants as being within a cluster if they were located within a certain number of base pairs of another significant GxE variant along the chromosome. For each variant type, we tested if the GxE variants were more clustered than expected by chance at a variety of cluster sizes/ using a permutation approach (**STAR Methods**). Upstream/promoter variants and synonymous variants were significantly more clustered than expected by chance at a range of cluster window sizes, while missense and downstream variants were not (**Fig, S4**). Interestingly, upstream/promoter variants were most significantly clustered at window sizes between 8 and 16 bp, which is similar to the size range of transcription factor binding sites (TFBS) in yeast. We chose to focus on a window size of 8 bp, which is roughly the average size of TFBS in yeast. At this distance, 36 out of 69 (52.2%) promoter GxE variants (targeting 34 unique genomic positions) were within a cluster, which is significantly more clustering of promoter GxE hit variants than expected by chance (permutation p = 0.002). The 36 clustered promoter GxE variants are located in 14 clusters, with all clusters sharing at least one significant pairwise GxE interaction at a relaxed threshold of FDR<0.1, though not necessarily in the same direction.

Interestingly, the *HMG1* promoter had three of these clusters, all of which had significant GxE interactions between caffeine and lovastatin, with strong effects on growth in lovastatin in both directions (**Figure 3h).** Interestingly, five of these clusters overlapped predicted transcription factor binding sites (TFBS) (Griffith et al., 2008; Harbison et al., 2004; Pachkov et al., 2013). For the other clusters, they may disrupt TFBS which have not been previously identified in the datasets we examined, perhaps due to context-specific binding, or may affect fitness through another mechanism.

### Gene-by-Environment interactions are Pervasive Among Natural Variants

Next, using our measurements of variant fitness effects in each condition, we examined how prevalent GxE interactions were among natural variants with significant fitness effects. If GxE interactions at the variant level were rare, we would expect to see that variants with strong fitness effects in one condition would mostly have similar effects in other conditions, and so would be correlated between conditions (**Figure 4a**).

**Figure 4:**
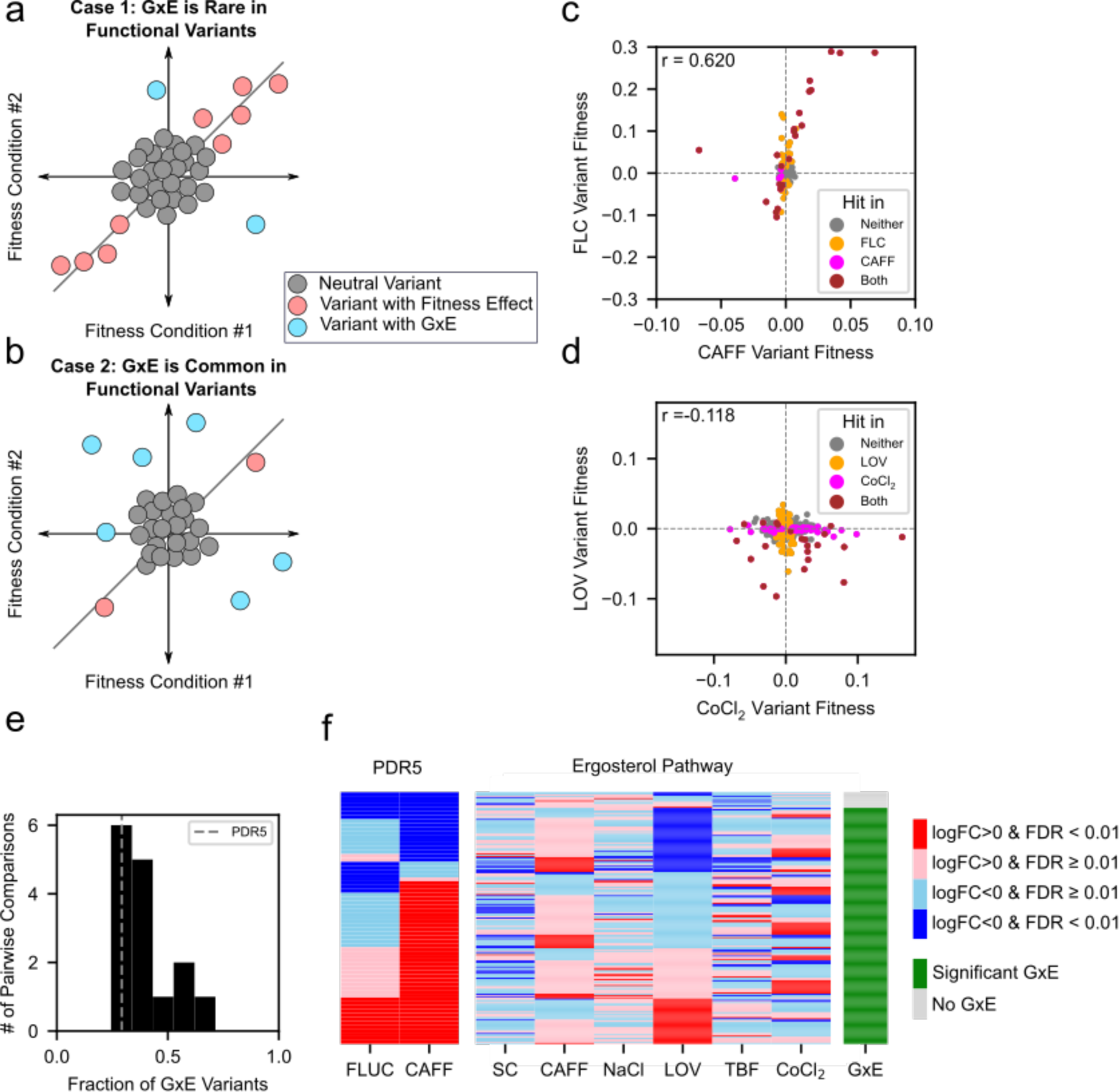
Quantifying GxE interactions among ergosterol pathway variants. *a,* Schematic of rare GxE between conditions (correlated effects). *b,* Schematic of common GxE between conditions (uncorrelated effects). *c,* Fitness effects of variants within *PDR5* in caffeine and fluconazole. *d,* Fitness effects of variants within ergosterol pool in lovastatin and CoCl2. *e,* Histogram showing the fraction of variants with significant fitness effects within a pair of conditions which show non-magnitude GxE for the ergosterol pool. *PDR5* variants measured in caffeine and fluconazole are shown as a dotted gray line. *f,* Heatmaps showing fitness effects of all variants with a significant effect in any condition. Significant positive effects (red), significant negative effects (blue), non- significant positive effects (pink), and non-significant negative effects (light blue). Far right column indicates variant with at least one GxE interaction (green) and without GxE (grey).

Conversely, if GxE interactions were common, we would see little correlation between fitness effects of the same variant in different conditions (**Figure 4b**). These patterns would only be visible for variants with measurable fitness effects, as those that are neutral or close enough to be not detected would be expected to show no correlation in either case. We saw examples of both of these patterns in our data, with fitness effects in caffeine and fluconazole within *PDR5* being generally well-correlated despite differences in magnitude, while ergosterol pathway variants measured in lovastatin and CoCl2 showed little to no agreement (**Figure 4c,d**).

Checking the fraction of GxE among hits for two conditions at a time, the fraction of variants with significant fitness effects in either condition with GxE interactions between the conditions ranges from 24.4 to 71.4% for the ergosterol pool conditions. *PDR5* variants in fluconazole and caffeine by this same method had 29.2% of significant variants showing GxE (**Figure 4e)**. Extending this analysis to examine effects in all conditions for the ergosterol pathway variants, it is clear that almost all variants with significant fitness effects showed GxE interactions (**Figure 4f)**. Among all variants measured in all six conditions which have at least one significant fitness effect in any condition, 93.7% have significant GxE interactions (Bootstrap 99% confidence interval 88.8%-97.6%). This result is robust to the FDR threshold used to define significant fitness effects and GxE interactions (**Figure S5**). In addition, this result is not solely influenced by a single condition (**Figure S6**). It’s important to note that having a strong fitness effect in one condition would make it more likely for a variant to have a detectable significant GxE interaction due to statistical power to detect a difference. However, if there existed a class of variants that showed consistent fitness effects across the conditions tested in magnitude and direction, they would have significant fitness effects while not showing GxE. In contrast, if all conditions had a similar fraction of GxE variants as the *PDR5* caffeine and fluconazole variants and were independent, only 82.2% (1-(1-.292)^5^) of variants would be expected to show GxE across six conditions. Strikingly, these analyses show that the vast majority of the non-neutral variants in the ergosterol biosynthesis pathway showed GxE, indicating that GxE interactions among natural variants are pervasive in this pathway.

### Regulatory GxE interactions in the ergosterol pathway

The finding that GxE interactions are pervasive among natural variants with detectable fitness effects in the ergosterol synthesis pathway led us to further investigate the pattern of their effects. In principle, the finding of pervasive GxE could be consistent with a scenario in which most variants have fitness effects in only one condition and are neutral in others (**Figure 5a**). In this case, we would expect to see that variants with a significant fitness effect in one condition were no more likely than any other variant to show a significant fitness effect in another, and so fitness effects across conditions should be distributed independently across the variants. Conversely, if variants with significant fitness effects in any condition were more likely to show strong fitness effects in other conditions (i.e. be pleiotropic), we would expect that the fitness distributions for the different conditions would not be independent, and significant fitness effects from these conditions would be more “clustered” in certain variants than expected by chance. 15.0% of variants measured in all six conditions showed a strong fitness effect in at least one condition, but 30.7% of variants that had a significant fitness effect in one condition had a significant fitness effect in at least one other condition, a two-fold enrichment (hypergeometric p = 5.27x10^-10^). These variants were further enriched for missense variants relative to all GxE variants (hypergeometric p = 5.40x10^-6^), and further depleted of synonymous variants (hypergeometric p = 1.82x10^-3^).

**Figure 5:**
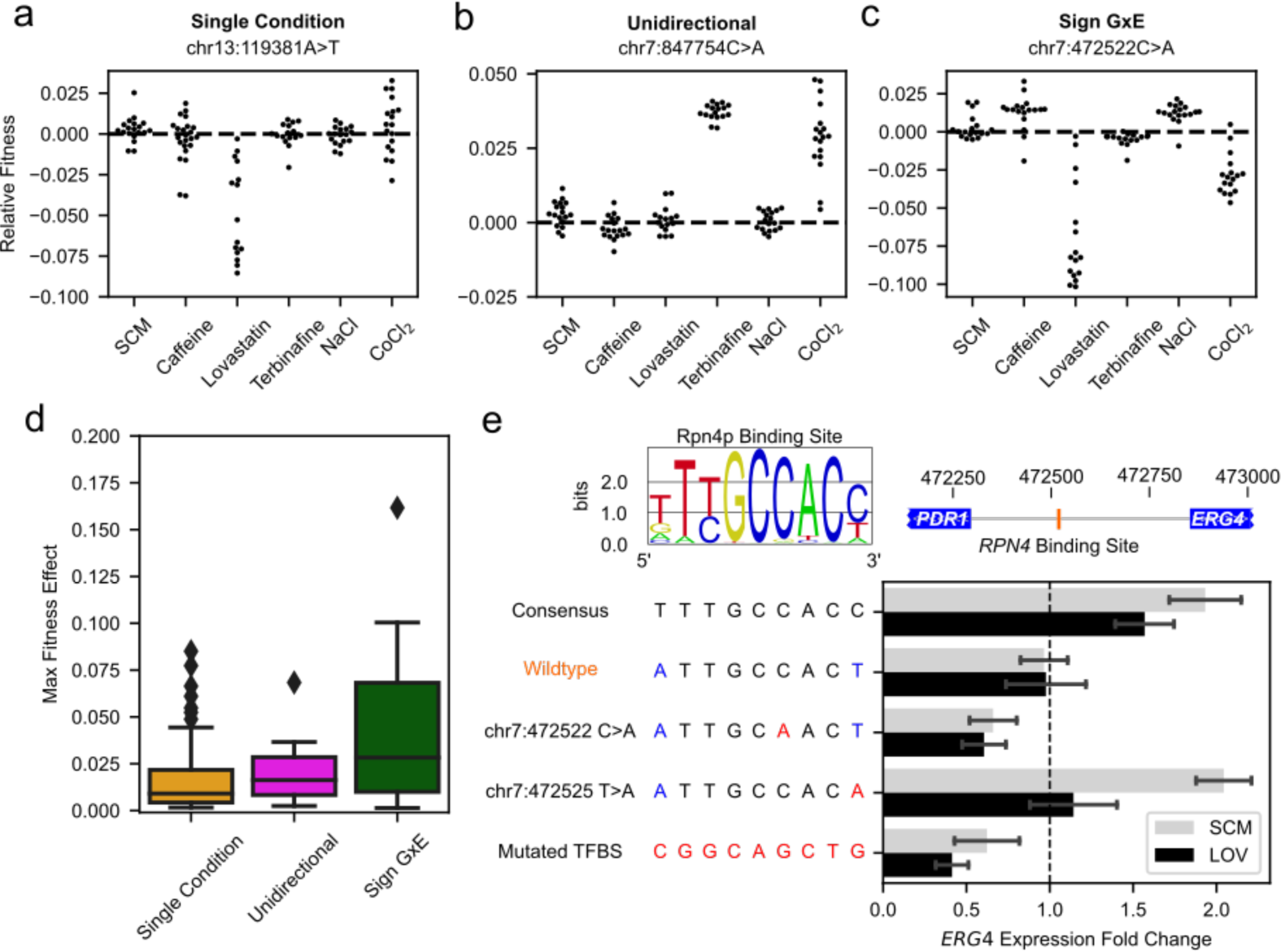
Types of GxE variants and effect of natural variation on *ERG4* expression. *a,* Example of fitness effect detected in only one condition. *b,* Example of fitness effects with same direction detected in two conditions. *c,* Example of fitness effects with opposite directions between conditions, showing sign GxE. *d,* Sign GxE variants have larger maximum fitness effects. Whiskers represent Q3 + 1.5xIQR and Q1 - 1.5xIQR, or the maximum and minimum values of the dataset if these are respectively lower or higher than the IQR based intervals. *e,* Effect of natural variants on *ERG4* expression. Top left: Consensus Rpn4p binding motif. Top right: Genomic location of Rpn4p binding site affected by chr7: 472522 C>A variant within *ERG4/PDR1* divergent promoter. Bottom left: Variants of Rpn4p binding site within *ERG4/PDR1* promoter tested. Bottom right: qRT-PCR measured expression of *ERG4* scaled by wildtype expression in an unedited strain, data presented as mean ± SEM.

Variants with significant effects in more than one condition can be grouped into two categories: 1) those with significant fitness effects in only one direction (**Figure 5b**) and those with significant fitness effects in opposite directions, which we term “sign GxE” (**Figure 5c**). Examining the strongest fitness effect for each of the variants with significant effects, we observed that variants showing sign GxE had significantly higher maximum effects than variants with significant effects in only one condition or multiple conditions in the same direction (Mann-Whitney U-test p=0.000193 and p =0.0367 respectively) (**Figure 5d**). This indicates that variants with more drastic effects on fitness in any given condition may be more likely to have fitness effects in the opposite direction in another condition.

In many cases, our single-nucleotide resolution suggests plausible molecular mechanisms underlying GxE. For example, we found that the pleiotropic variant exhibiting sign GxE at chr7: 472522 C>A was located in a canonical Rpn4p binding site (**Figure 5e**) (Foat et al., 2008; Mannhaupt et al., 1999). This variant’s strongest effect was a significant fitness decrease in lovastatin. Since Rpn4p is a transcriptional activator, we hypothesized that the disruption of the Rpn4p binding site might decrease *ERG4* expression. We used RT-qPCR to measure expression of *ERG4* in a genotyped strain carrying chr7: 472522 C>A and found that its expression decreased relative to the wildtype strain (**Figure 5e**). This decrease in expression agreed with *ERG4* expression in a strain carrying a fully ablated Rpn4p TFBS, while a strain carrying an Rpn4p consensus site had higher *ERG4* expression. Interestingly, a strain with another natural variant that mutated a lower information base within the Rpn4p binding motif (chr7: 472525 T>A) showed a slight fitness decrease in lovastatin and did not show a significant decrease in *ERG4* expression. Therefore, we reasoned that the chr7: 472522 C>A variant disrupted *ERG4* expression through mutation of the Rpn4p TFBS. We then further tested the fitness in the Rpn4p consensus and Rpn4p mutated TFBS, showing that *ERG4* expression correlated with fitness in lovastatin (**Figure S7**). In sum, we showed that CRISPEY-BAR was able to survey thousands of natural variants and identify the variants affecting fitness at the nucleotide-level, directly leading to discovery of molecular mechanisms of GxE interactions.

## Discussion

We demonstrated that the CRISPEY-BAR strategy and its applications provide a solution to rapidly discover natural genetic variants impacting a complex trait. As a proof of principle, we pinpointed 548 variants with significant effects on growth within QTLs, as well as across a core metabolic pathway. Although we potentially expected QTLs to contain variants with GxE interactions, it is surprising to uncover hidden complexity among natural variants in close proximity to each other, which harbor fitness effects in opposite directions (**Figure 2**). Within the ergosterol synthesis pathway, we were not only able to find natural variants that facilitate differential drug responses, but also reveal the pervasiveness of GxE interactions among variants with fitness effects across diverse environmental challenges (**Figures 3, 4**). In future studies it will be interesting to see if the pervasive GxE interactions we observed for the ergosterol pathway will apply to additional pathways, growth conditions, and species.

We have demonstrated that pooled fitness measurements are reproducible across CRISPEY-BAR experiments with only minimally overlapping sets of variants (**Figure 1j**). This indicates that independent CRISPEY-BAR experiments with mostly separate sets of variants may be directly compared by designing overlapping variants as fitness standards between pools for joint analysis, allowing us to scale CRISPEY-BAR experiments to explore a greater number of natural variants. Scaling up to cover variants across entire genomes, we will be able to carry out even deeper probing of the relationship between genotype and any trait amenable to pooled phenotyping (including any traits that can be tied to growth or fluorescence-based reporters).

Deciphering the non-coding genome has been a major challenge even in just one experimental condition and is further complicated by GxE interactions. In this study, we showed that a class of variants with GxE cluster tightly within promoter regions, and further found that some of them overlap known TFBS (**Figure 3g**). Although GxE variants are most highly enriched in missense variants, we found no genomic clustering of these protein-altering variants; elucidating their molecular mechanisms will be an exciting area for future study.

CRISPEY-BAR is highly efficient in precise editing, and we envision multiple routes to further improve its effectiveness. The RT was shown in CRISPEY to be effective in production of msDNA as DNA donors for precision editing (Sharon et al., 2018). We have since tested additional retron RTs in CRISPEY, showing higher efficiency in yeast, as well as editing activity in human cells (Zhao et al., 2022). This study utilized the SpCas9 with an ‘NGG’ PAM site, limiting the variants that can be targeted; however, alternative nucleases with different PAM sequences can be interchanged with SpCas9 to target additional variants (Hu et al., 2018; Legut et al., 2020; Nishimasu et al., 2018).

The CRISPEY-BAR approach has an efficient guide for barcoding, while the variant editing guide can have a range of efficiency. Because we deployed two or more untested guides to target each variant, we are more likely to believe that the guides that show the same significant fitness effect were both efficient in making precise edits. Moreover, the six UMIs allow outlier detection where spontaneous mutations or off-target effects may have taken place. Combining guide reproducibility and UMI editing-competition replication, every CRISPEY-BAR experiment allows us to accumulate data points for supervised learning of effective guide design in CRISPEY-based editing strategies.

In this study, CRISPEY-BAR was applied to a lab strain of budding yeast to evaluate the effect of natural variants. This may limit the portability of the fitness effects we measure for individual variants, since they are only measured in this lab strain genetic background. This caveat can be overcome by applying CRISPEY-BAR to additional strains of budding yeast to not only capture the effects of variants within one lab strain, but also the effect of genetic background (see companion paper by Ang *et al*.). Looking ahead, the CRISPEY-BAR design also allows for additional ribozymes and CRISPEY cassettes to be incorporated. A single barcode-insertion cassette plus two or more variant editing cassettes can be expressed in the same transcript, allowing simultaneous editing of two genetic variants of choice and integration of a variant-pair specific barcode.

With this design, we will be able to observe gene-by-gene (epistatic) interactions, as well as gene-by-gene-by-environment (GxGxE) interactions that govern the crosstalk between gene networks and the environment (Costanzo et al., 2016, 2021; Jaffe et al., 2019).

The observation that GxE interactions were found to be pervasive among variants with fitness effects in just six conditions tested was a surprising result. Most of the variants with GxE have a significant effect in only one condition, which by definition shows GxE with respect to the rest of the conditions. More excitingly, we found a fraction of the variants to harbor sign GxE, which implies fitness tradeoffs in fluctuating environments where selection acts in opposite directions on the variant (**Figure 4g**). Moreover, we found a trend in which large-effect variants tend to also have larger effects in another condition than expected by chance, forming a class of pleiotropic variants with two or more conditional effects. While we expect additional variants with fitness effect to be identified as more conditions or drug conditions are tested on the same set of variants, it is intriguing to think that the pleiotropic variants may harbor disproportionate amounts of environment-specific effects. If such is the case, by performing a limited set of CRISPEY-BAR experiments with a diverse set of conditions, we will be able to prioritize a set of pleiotropic variants that are likely to have effects in the remaining, untested conditions spanning the phenotypic space.

## Acknowledgments

We thank M.M. Desai, D. Petrov and members of the Fraser lab for helpful discussions. S.A.C. was partially supported by Bio-X Stanford Interdisciplinary Graduate Fellowship, NIH grant 1F31ES030282 and NIH 5T32GM007276-42. A.F.K. was partially supported by NIH NHGRI Stanford Genomic Training Program 5T32HG000044. R.M.L.A. was partially supported by the National Science Scholarship, from the Agency of Science, Technology and Research (A*STAR). This work was supported by NIH grants 2R01GM097171, 1R01GM134228 and 1F31ES030282.

## Author Contributions

Conceptualization, H.B.F., S.A.C and A.F.K.; Investigation, S.A.C., A.F.K., Y.X and H.B.F.; Validation, S.A.C. and A.F.K.; Formal Analysis, S.A.C. and A.F.K.; Data curation, S.A.C. and A.F.K.; Methodology, Software and Visualization, S.A.C., A.F.K. and R.M.L.A.; Writing – Original Draft, S.A.C, A.F.K, H.B.F; Writing – Review & Editing, S.A.C, A.F.K, R.M.L.A. and H.B.F; Funding Acquisition: S.A.C. and H.B.F.; Supervision H.B.F..

## Supplemental figures

**Supplemental Figure S1:**
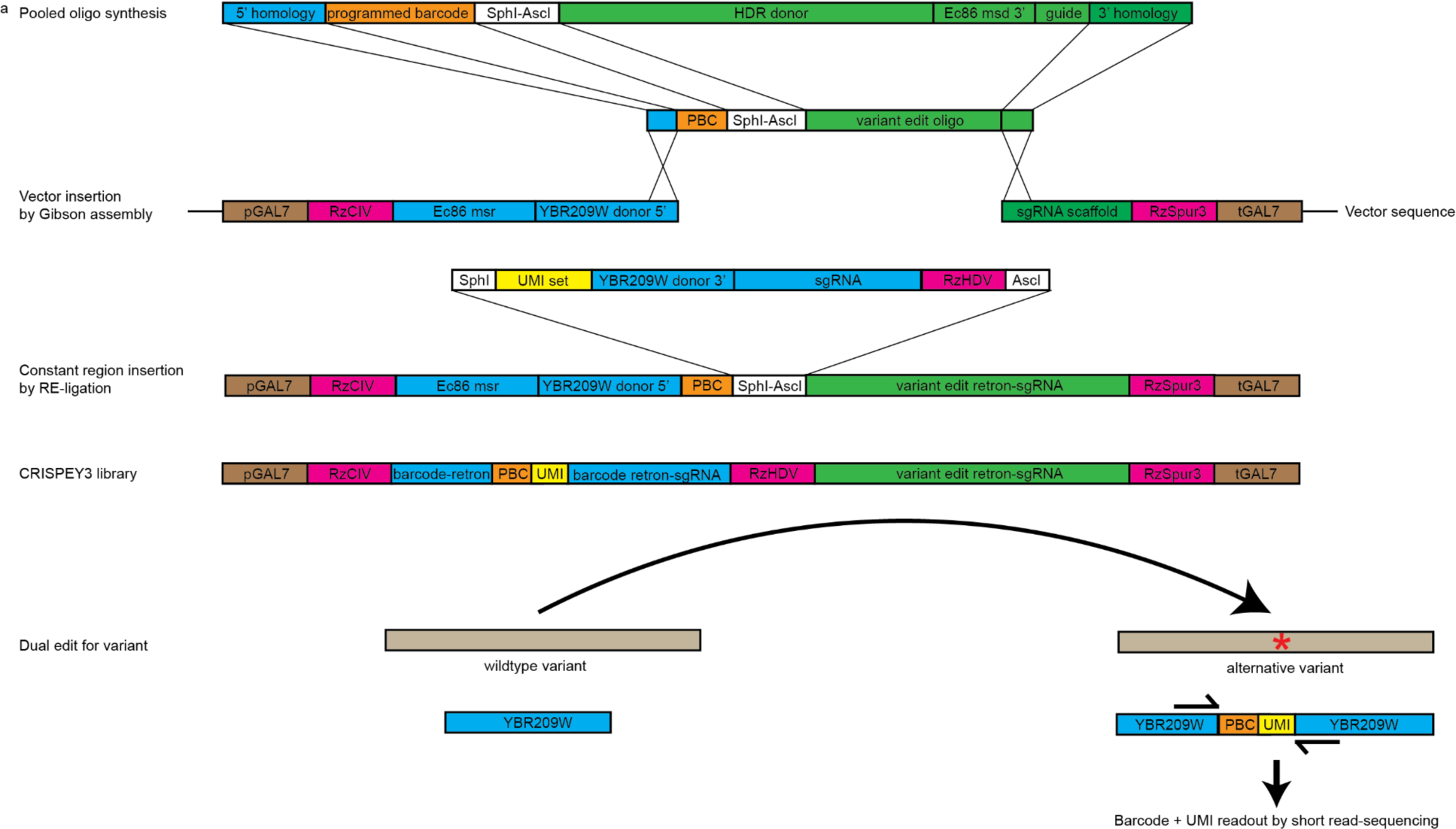
Schematic for library cloning in CRISPEY-BAR. Pooled oligonucleotide libraries, empty vector sequence (pSAC200) and an example of ligated *ADE2* editing vector (pSAC212) can be found in Supplementary Table S1.

**Supplemental Figure S2:**
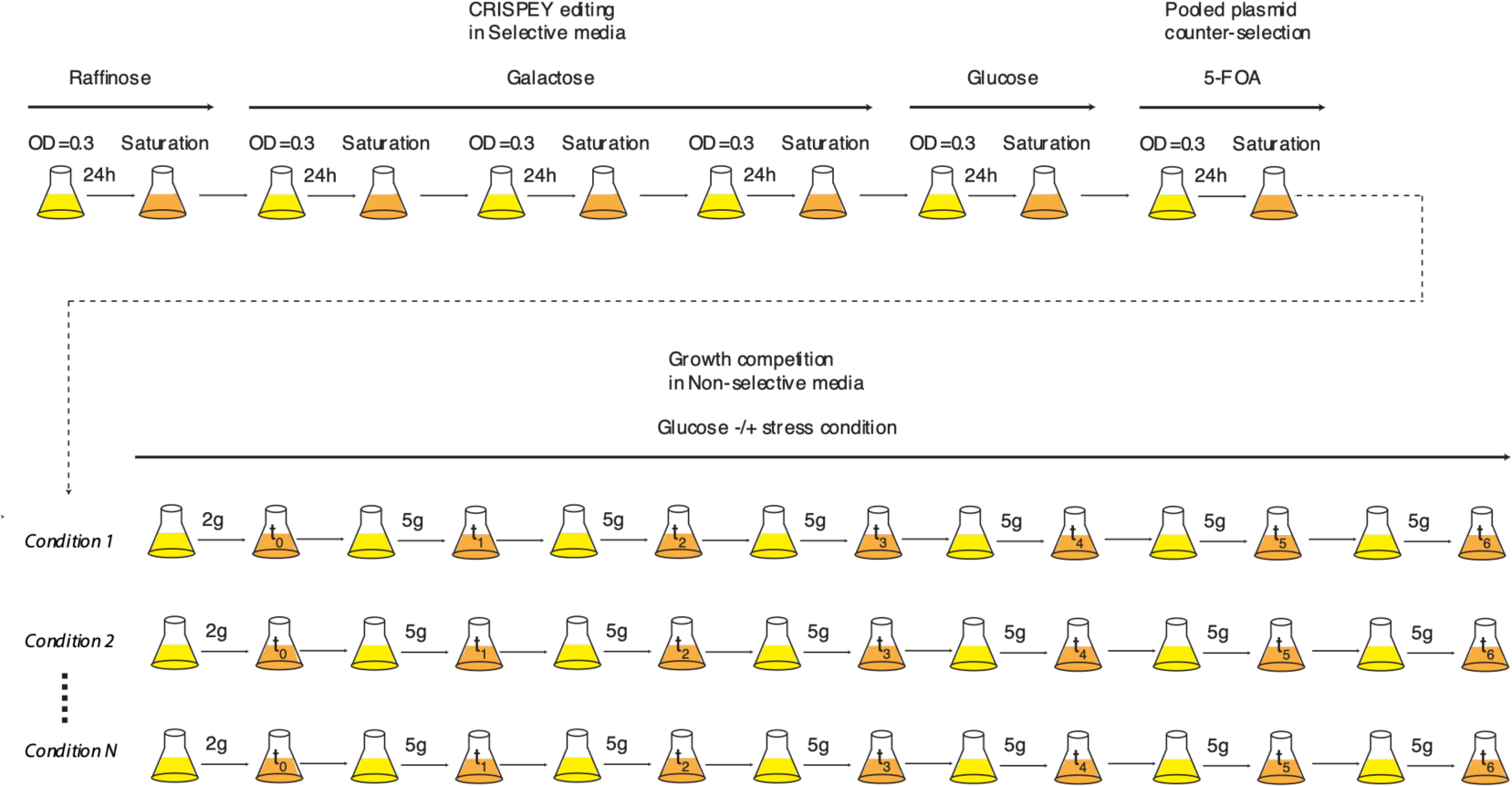
Schematic for pooled editing and growth competition in CRISPEY-BAR. STAR Methods for detailed description.

**Supplemental Figure S3:**
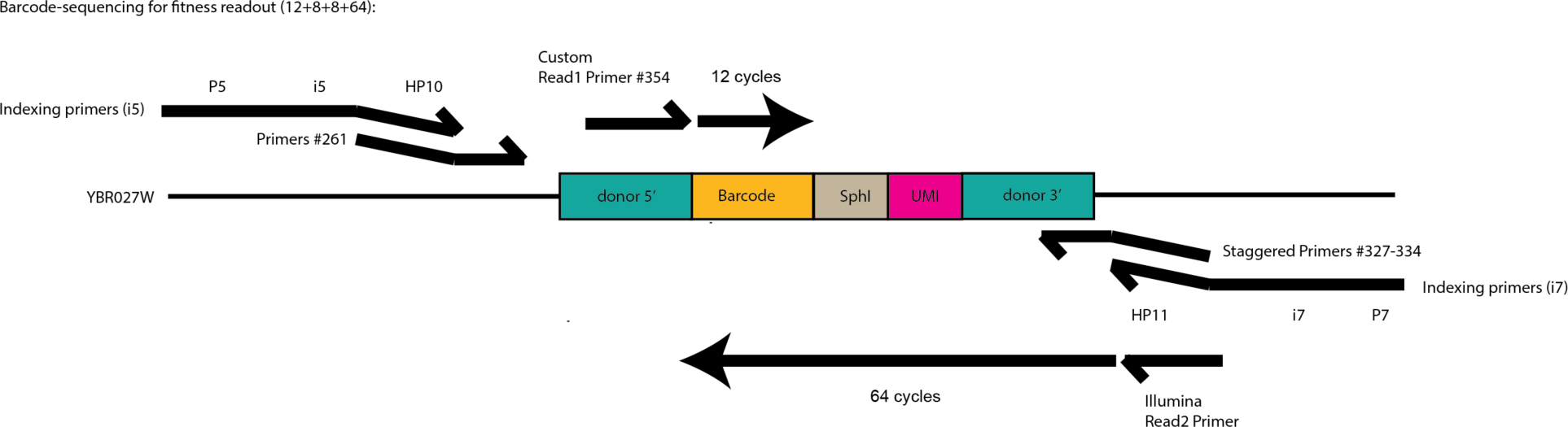
Schematic for CRISPEY-BAR sequencing library preparation. Primer sequences can be found in Supplementary table S1.

**Supplemental Figure S4:**
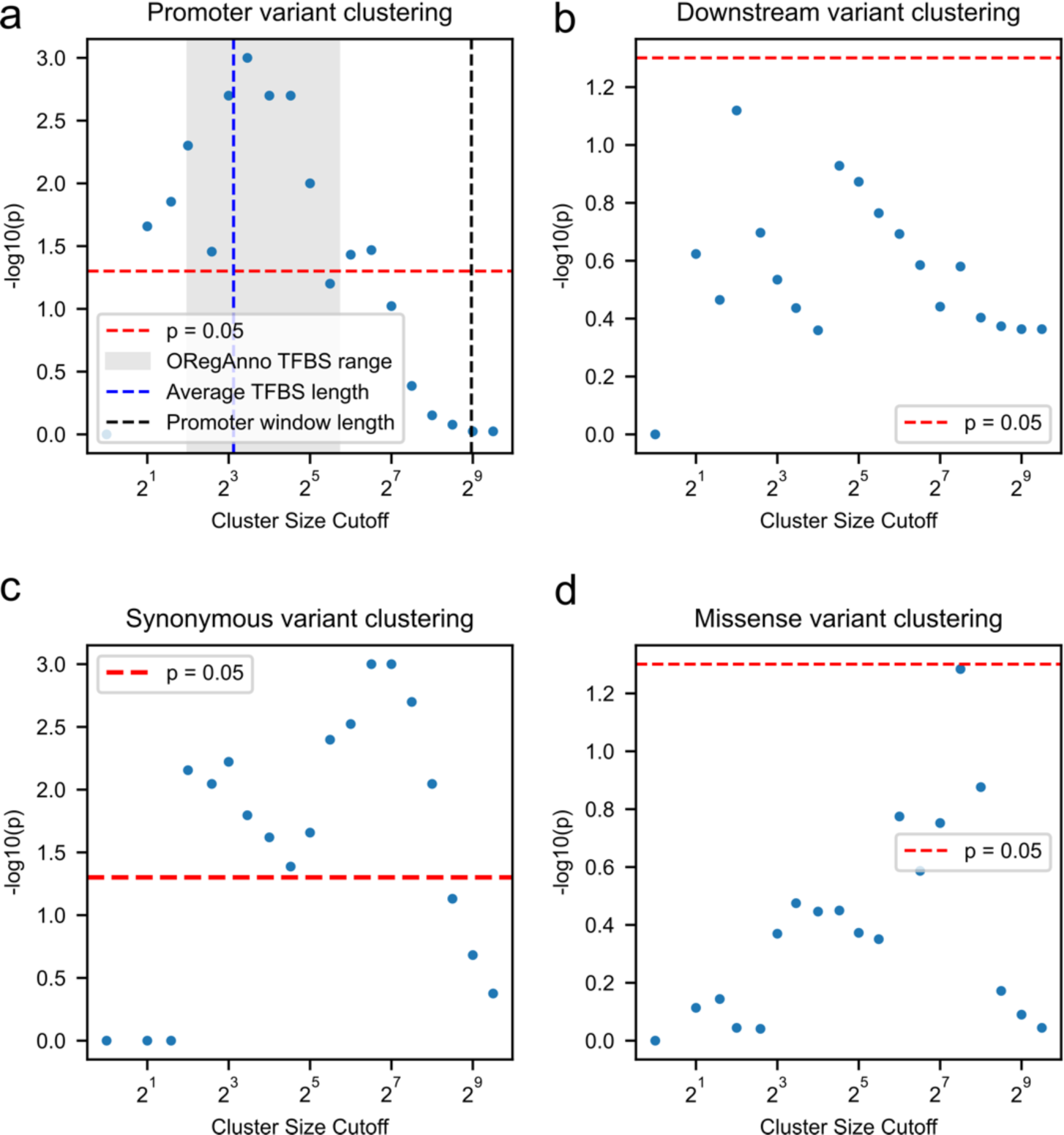
Clustering analysis for different variant types. a, Promoter variant clustering. b, Downstream variant clustering. c, Synonymous variant clustering. d, Missense variant clustering.

**Supplemental Figure S5:**
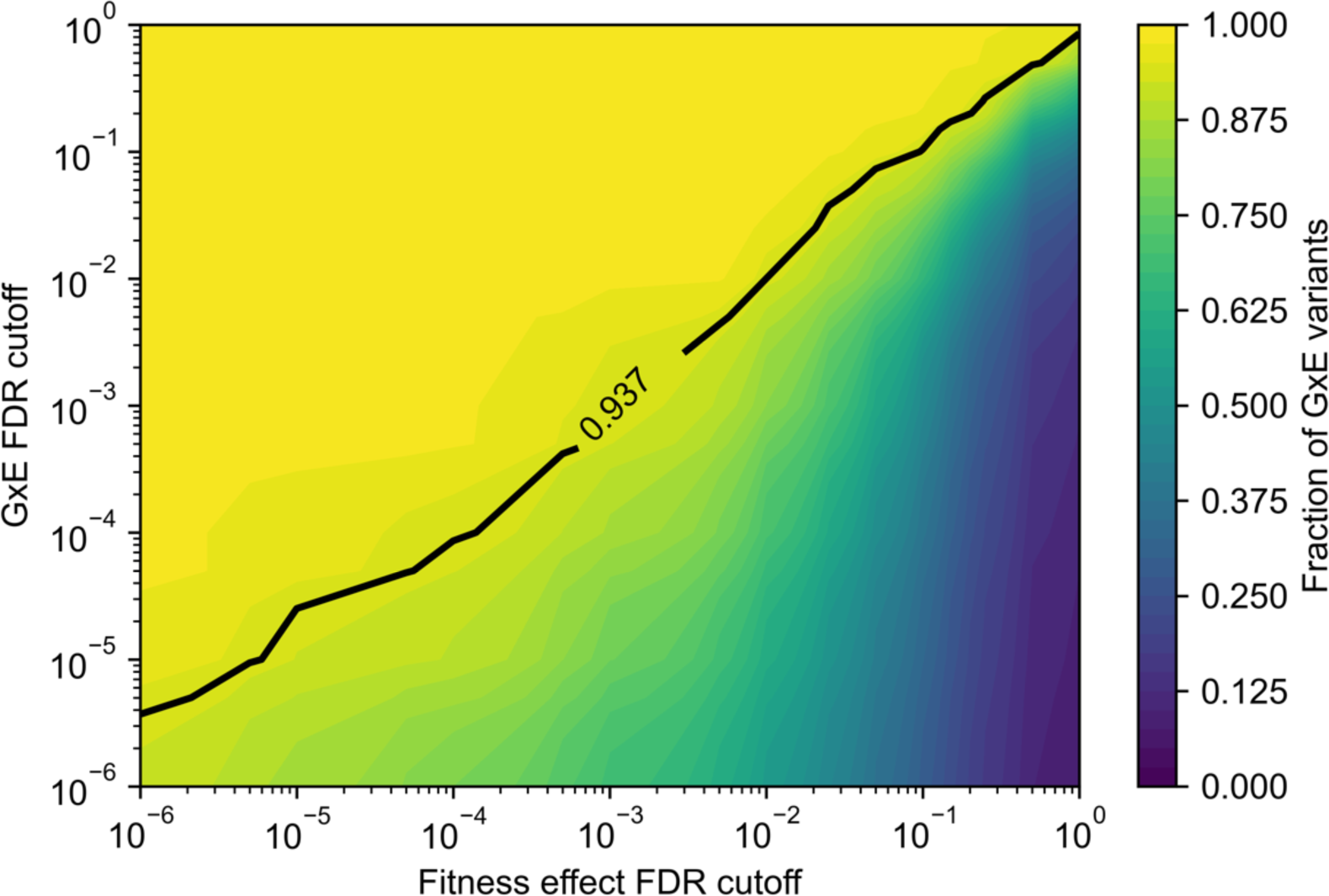
Overlap between variants with effects on fitness and GxE variants at differing FDR thresholds. Line indicates measured overlap at FDR<0.01 for both thresholds for variants measured in all six conditions.

**Supplemental Figure S6:**
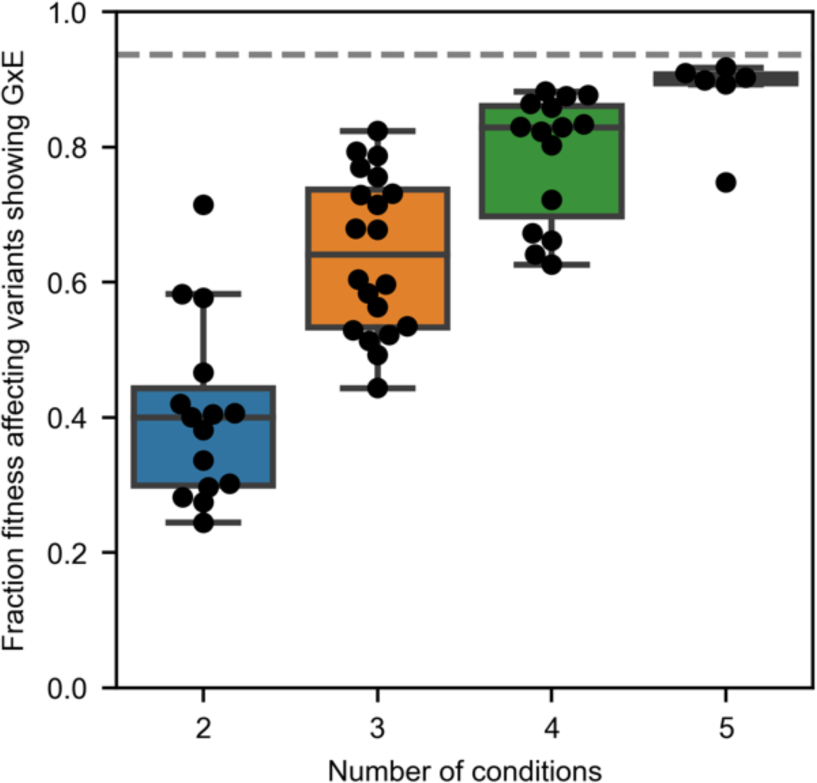
Fraction of variants with fitness effects harboring GxE is not dependent upon specific conditions analyzed. Each dot indicates fraction of variants with fitness effects harboring GxE across conditions analyzed, where the conditions are subsets of the 6 conditions in Figure 3d. Dashed line indicates fraction of GxE across all six conditions.

**Supplemental Figure S7:**
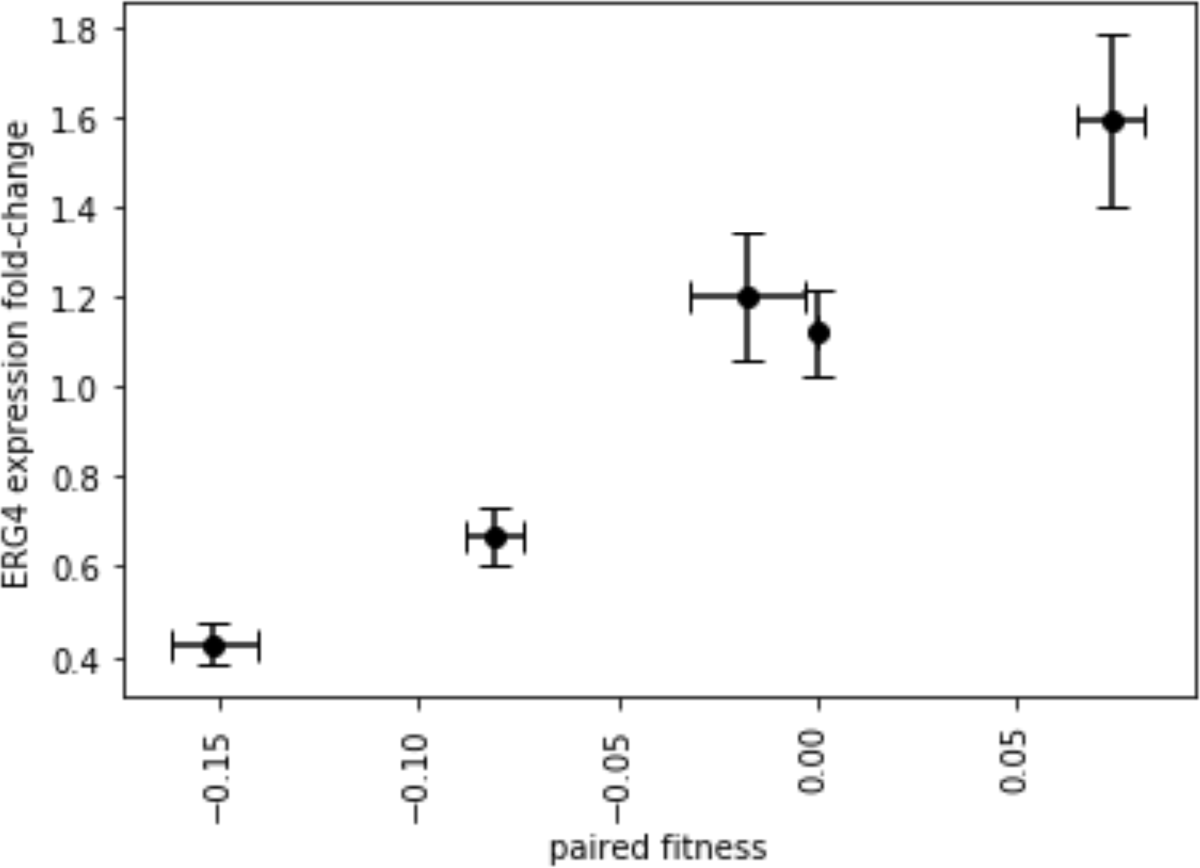
Fitness and *ERG4* expression for variants in Figure 5e. X-axis: Paired fitness from flow cytometry measurements similar to Figure 1i, see also **STAR Methods**. Y-axis: *ERG4* expression change same as shown in Figure 5e. Data presented as mean ± SEM.

## STAR ★ Methods

### Contact for Reagent and Resource Sharing

Further information and requests for reagents may be directed to and will be fulfilled by the corresponding author Hunter Fraser.

### Experimental Model and Subject Details

All strains used in this study were derivatives of S. cerevisiae BY4742 (Brachmann et al., 1998). Construction of strains with integrated SpCas9 and Ec86 Reverse Transcriptase was described previously (Sharon et al., 2018).

### Method details

#### Variant selection and pooled oligonucleotide design

Natural variants were sourced from the 1,011 genomes project documented the following criteria (Peter et al., 2018). For QTL fine mapping, QTLs (Bloom *et al*., 2019 supplementary file elife-49212-Figure3-data1-v2.xls, sheet_name=’within-cross model’) were filtered for QTLs containing only one gene, have a q-value > 0.05, then ranked by Beta_abs for maximum effect size (Bloom et al., 2019). We excluded the following genes to avoid interference with CRISPEY editing and genes unavailable from the base strain genotype: HO, HIS3, URA3, LEU2, LYS2, GAL1, GAL3, GAL4, GAL7, GAL10, GAL80, HAP1 and POLR2. The QTL borders were defined by coordinates within ’1.5 LOD drop CI, left’ and ‘1.5 LOD drop CI, right’ as annotated in Bloom, 2019, and gene regions were defined by +-500bp from the coding region (Bloom et al., 2019). Natural variants within the union of the QTL borders and the gene region were included in the library corresponding to the traits, excluding singletons and doubletons (Peter et al., 2018). The traits include growth in: ’Cobalt_Chloride;2mM;2’, ’Caffeine;15mM;2’ and ’Fluconazole;100uM;2’, and we refer to these traits as ‘stress conditions’ (Bloom et al., 2019). For the ergosterol pool, all non-reference alleles from yeast natural variants that were within +-500bp from the coding region of the selected ergosterol pathway genes were included^4^. We targeted more than 1000 variants per QTL condition pool, and all possible variants for the ergosterol pool (**Figure 2a,3a**). We designed CRISPEY oligos to edit these variants in the ZRS111 strain, which contains the S288c reference alleles. The guides and donors selected for CRISPEY editing were designed as described, with the following parameters or modifications (Sharon et al., 2018): 1. The alternative allele is within -6 to -1 and +1 to +2 positions of the guide target and PAM sequences; 2. The donor template is 108 bp in length with asymmetric homology arms, 40 bp for the 5’ arm and 68 bp for the 3’ arm; 3. Variants were included if two or more guides were found for a given variant. The resulting msDNA donor will result in a shorter 3’ homology arm and longer 5’ arm flanking the variant, which was to have higher HDR efficiency using ssDNA as repair donor (Richardson et al., 2016). The donors were further filtered to exclude SphI, AscI and NotI restriction sites used in the cloning process, as well as keeping a minimum of 30 bp homology arm 5’ of variant and 55 bp 3’ of homology arm in the donor template. The resulting output is 250 bp per oligo, consists of 5’ homology to the pSAC200 CRISPEY-BAR vector, 12 bp programmed barcode, restriction site region for cloning, 108 bp donor template sequence, 34 bp constant region, 20 bp guide sequence and 3’ homology to the pSAC200 CRISPEY-BAR vector (**Supplemental Figure 1**).

Specifically, the general sequence is: 5’- GTTGCAGTTAGCTAACAGGCCATGCNNNNNNNNNNNNGCATGCAGCGGCCGCAGGCGCGCCNNNNNNNNNNNNNNNNNNNNNNNNNNNNNNNNNNNNNNNNNNNNNNNNNNNNNNNNNNNNNNNNNNNNNNNNNNNNNNNNNNNNNNNNNNNNNNNNNNNNNNNNNNNNAGGAAACCCGTTTCTTCTGACGTAAGGGTGCGCANNNNNNNNNNNNNNNNNNNNGTTTCAGAGCTATGCTGGAAACAGCAT-3’, where the first 12 Ns represent programmed barcodes, the following 108 Ns represent donor template sequence, and the last 20 Ns represent guide sequence.

### Programmed Barcode Design

Barcodes were designed using a custom script implementing a quaternary Hamming(12,8) code based on the encoding scheme described in a previous study (Bystrykh, 2012). This encoding scheme generates DNA barcodes with a minimum Hamming distance of 3, allowing for error correction of 1 bp mutations or DNA sequencing errors. The list of all Hamming(12,8) DNA barcodes was then filtered to remove barcodes containing restriction sites used in our cloning process, Illumina i5 and i7 Nextera handles, homonucleotide stretches greater than length 3, dinucleotide repeats greater than length 5, any 12 bp section of pSAC200, and any 12 bp section of our custom sequencing primers. In addition, Primer3 was used to predict any hairpin structures, and if a structure was found, that barcode was removed (Koressaar and Remm, 2007). The final list of barcodes was then assigned to 392 possible wells, ensuring that barcodes within each well had a minimum Hamming distance of 5, theoretically enabling error correction of sequencing errors in up to 2 bp for barcodes within the same well.

### Library cloning

Primers and additional oligonucleotides can be found in **Supplemental Table S1.** Oligonucleotide (Twist Biosciences) libraries were ordered in the format of 192 wells, each well containing 121 oligonucleotides each. This format allows pooling of oligonucleotides in combinations relevant to each competition experiment. Each well included 119 variant editing oligonucleotides, 1 control oligonucleotide with a non-editing guide (sgGFP) and 1 control oligonucleotide editing a 8-bp frameshift deletion as a positive control with gene knockout effects adapted from a previous study(Bao et al., 2018).

Oligonucleotides were first amplified with Q5 polymerase (NEB) with 1 uM primer #615 and #576 in 50 uL reaction following manufacturer instructions and initial denaturation of 98°C for 2 min, and then 5 cycles of 98°C for 10 s and 65°C for 30 s, followed by 25 cycles of 98°C for 10 s and 69°C for 40 s, then final extension of 72°C for 2 min. PCR products were then purified with 45 uL nucleoMAG NGS beads (hereafter, “beads”) (Takara) and eluted with 20 uL water. 2 uL of the first round PCR product was further amplified with Q5 polymerase (NEB) with 1 uM primer #615 and #576 in 50 uL reaction as manufacturer instructions and initial denaturation of 98°C for 2 min, and then 15 cycles of 98°C for 10 s and 69°C for 30 s, then final extension of 72°C for 2 min. Second round PCR products were then purified with 45 uL beads and eluted with 20 uL Tris pH 8.0. We quantified and pooled PCR products from each well by equal volume to the assigned pools (**Supplemental Table S2**).

The pooled oligonucleotides PCR products were purified using SizeSelect II 2% gel (Invitrogen), followed by bead purification and prepared NGS libraries to quantify the counts from each well. Briefly, the pooled oligos were amplified with Q5 polymerase (NEB) with 1 uM primer #617 and #337-343 in 50 uL reaction following manufacturer instructions and initial denaturation of 98°C for 2 min, and then 15 cycles of 98°C for 10 s and 69°C for 40 s, then final extension of 72°C for 2 min, followed by purification using 45 uL beads and indexing PCR using Illumina dual-indexing primers. The indexed amplicons corresponding to each pool were then sequenced by MiSeq using reagent kit v2 Nano to obtain paired-end 150bp reads that are mapped to the designed oligonucleotides. We counted the relative proportions of oligonucleotides from each well in the assigned pool, then re-pooled the PCR products again with normalized volumes to target equal molarity between wells in each pool.

The pSAC200 empty vector was digested twice with NotI-HF (NEB) and Quick CIP (NEB), and the linearized vector was purified using beads. 290 ng of linearized pSAC200 vector and 140 ng of well-normalized, pooled oligonucleotide PCR products from above were assembled in 20 uL NEBuilder HiFi mastermix (NEB) reaction according to manufacturer instructions, with 1:10 molar ratio between vector:insert. The assembled products were purified by beads and eluted in 10 uL water. 3 uL of the assembled products were used for electroporation with 27 uL Endura Electrocompetent cells for CRISPR DUO (Lucigen). Two electroporation reactions were performed for each pool following manufacturer instructions and recovered in SOC media (Lucigen) for 25 min at 37°C and plated to a single 15 cm LB agar plate with Carbenicillin (GoldBio). A serial dilution of the recovered bateria was plated to estimate colony forming units (cfu), and all pools contained more than 500,000 cfus. The transformants were incubated for 22 hr at 32°C and the resulting bacterial lawn was collected for storage in LB with 10% glycerol at -80°C. Half of the collected transformant stock was used for plasmid extraction using Nucleobond Xtra Midi Plus (Macherey-Nagel) and eluted as “post-Gibson” plasmid pools, yielding 105-120 ug of plasmid DNA.

20 ug of post-Gibson plasmid pools were digested twice with SphI-HF(NEB), AscI(NEB), Quick-CIP(NEB) and NotI-HF (NEB), purified by beads and eluted in 12 uL 10mM Tris 8.0 as ligation vectors. A mixture of six UMI associated ligation inserts was generated by six 100 uL reactions Q5 (NEB) PCR reaction with one of six forward primers: #591, #592, #594, $506, #603 and #604; and reverse primer #590, with plasmid pSAC212 as template. PCR was performed with 1 uM of each primer as manufacturer instructions, and initial denaturation of 98°C for 3 min, and then 35 cycles of 98°C for 10 s, 66°C for 30 s, 72°C for 40 s; then final extension of 72°C for 2 min. The ligation insert PCR products were digested with SphI-HF (NEB) and AscI (NEB) and bead purified, then pooled in equal molar into a mixture of six UMI ligation inserts. 1 ug of the linearized pool vectors were ligated to 1.5 ug of six UMI ligation mix (vector:insert=1:30) with 10 uL T4 ligase (NEB) in 100 uL 1x T4 ligase buffer at 16°C overnight.

The ligation product was purified by beads and eluted in 30 uL water. 3 uL of the purified ligation products were used for electroporation with 27 uL Endura Electrocompetent cells for CRISPR DUO (Lucigen). Two electroporation reactions were performed for each pool, one reaction with ligation insert and the other without insert as negative control.

Electroporation was performed following manufacturer instructions and recovered in SOC media (Lucigen) for 30 min at 37°C and the with-insert ligations were plated to two 15 cm LB agar plates with Carbenicillin (GoldBio) at 32°C for 22 hr. A serial dilution of the recovered bacteria from both with- and without-insert ligations was plated to estimate cfu, and all pools contained more than 1,000,000 cfu, corresponding to at least 2,500x coverage for each oligonucleotide on average within each pool. Ligation plates were incubated at 32°C for 22 hr, and transformants were stored in LB with 10% glycerol.

Ligated plasmids were extracted from one fourth of the collected bacteria from each pool using Nucleobond Xtra Midi Plus (Macherey-Nagel) and eluted as “post-ligation” plasmid pools, yielding 160-240 ug of plasmid DNA per reaction.

### Yeast transformation, editing induction and plasmid curing

The base strain ZRS111 was described previously(Sharon et al., 2018). 4 ug of the post- ligation plasmid pools were digested with NotI-HF (NEB) and quick-CIP(NEB) and directly transformed into the yeast strain ZRS111 by LiOAc heat shock transformation (Gietz and Schiestl, 2007). The yeast transformant pools were selected on YNB - histidine -uracil 2% glucose (1.7g/L yeast nitrogen base (RPI); 5 g/L Ammonium Sulfate (ACROS organics); 1.9 g Dropout synthetic mix minus histidine, uracil w/o nitrogen base (US Biological) and 20 g/L glucose (Sigma) 2% agar plates and stored in YNB -histidine - uracil 2% glucose media with 15% glycerol at -80°C. Yeast transformants containing post-ligation pools were inoculated to 200 mL YNB -histidine -uracil 2% raffinose (1.7 g/L yeast nitrogen base (RPI); 5 g/L Ammonium Sulfate (ACROS organics); 1.9 g Dropout synthetic mix minus histidine, uracil w/o nitrogen base (US Biological) and 20 g/L raffinose (Sigma) media starting at OD600=0.4, shaking at 30°C for 16 hr (**Supplemental Figure S2**). The raffinose cultures were further re-inoculated in 200 mL YNB -histidine - uracil 2% galactose media starting at OD600=0.4 and shaking at 30°C for 24 hr three times, for a total of 72 hr in galactose media in order to induce CRISPEY-BAR editing.

Cells were harvested from the last galactose media growth and stored in YNB -histidine - uracil 2% glucose media with 15% glycerol at -80°C. Edited cells were then plasmid-cured by growing in 200 mL YNB 2% glucose (1.7g/L yeast nitrogen base (RPI); 5 g/L Ammonium Sulfate (ACROS organics); 1.9 g Dropout synthetic mix complete, w/o nitrogen base (US Biological) and 20g/L glucose (Sigma) media starting at OD600=0.4, shaking at 30°C for 16 hr, then re-inoculated to YNB 2% glucose media with 1 g/L 5- Fluororotic acid monohydrate (GoldBio) starting at OD600=0.4, and shaking at 30°C for 24 hr(Boeke et al., 1987). The plasmid-cured cells were collected and stored in YNB 2% glucose media with 15% glycerol at -80°C.

### Pooled competition

Pooled competitions were carried out in 1 L baffled flasks in YNB 2% glucose (SC, hereafter) media with or without specified conditions (**Supplemental Figure S2**). For stress conditions, we used the following final concentrations in SC for the stress reagents: sodium chloride (0.8M); fluconazole (7.5 ug/mL); cobalt chloride (1.5 mM); terbinafine (40 ug/mL); lovastatin (30 ug/mL, stock solution was dissolved in 15%(v/v) ethanol followed by heat activation); caffeine (1 mg/mL). The concentration of each drug/salt was titrated to approximately 5 generations of growth of the ZRS111 strain every 12 hr, indicating overall decreased fitness in each condition to apply consistent growth stress to cells. In contrast, for SC media only, there are approximately 5 generations of growth ZRS111 strain in 8 hr. Cells were thawed in 200 mL SC media from glycerol stock starting at OD600=0.4 and grown at 30°C shaking at 250 RPM. Cells were passaged every 12 hr and diluted to fresh 1 mL SC media with specified conditions, and every 8 hr for SC media only. Five intervals separated by six timepoints (T1∼T6) were harvested at every time point once passage was complete. Harvested cells were spun down, washed with water, and stored at -20°C.

### Sequencing library preparation

Yeast genomic DNA was extracted from 60 - 80 OD of each sample using the MasterPure Yeast DNA Purification Kit (Lucigen) with four reactions per sample. Genomic DNA was eluted in 200 uL per sample, further digested with 1 uL RNaseA and quantified by Qubit dsDNA HS assay (Invitrogen). 10 ug of genomic DNA was amplified in 400 uL Q5 polymerase (NEB) PCR reaction with 1 uM forward primer #261 and 1 uM reverse primer equimolar mix of primers #327∼#334 (**Supplemental Figure S3**). PCR was performed following manufacturer’s instructions, with 1M Betaine and initial denaturation of 98°C for 2 min, then 19 cycles of 98°C for 10 s, 65°C for 20 s; then extension at 72°C for 5 min. 100 uL of first round of PCR products were purified using 100 uL beads and 15 uL of the purified amplicons were further indexed by 50 uL Q5 polymerase (NEB) PCR reaction following manufacturer’s instructions with 1 uM equimolar mix of indexing primers for Illumina sequencing, and initial denaturation of 98°C for 2 min, then 8 cycles of 98°C for 10 s, 70°C for 20 s; then extension at 72°C for 2 min. The indexed amplicons were purified with 50 uL beads, eluted in 100 uL water and quantified by Qubit dsDNA HS assay (Invitrogen). The purified, indexed amplicons from six time point samples for the three replicates per competition were mixed equimolar and purified by SizeSelect II gel (Invitrogen) for ∼300 bp product. The size selected libraries were then purified by beads and submitted for paired-end sequencing on NextSeq 550 using custom read1 primer #354, with custom cycles of 12 cycles for read1, 8 + 8 cycles for dual indices and 64 cycles for read2 using a 1 x 75 bp High-Output Kit (**Supplemental Figure S3**).

### Read Processing

Reads in fastq format from competition libraries sequenced using NextSeq were processed using a custom script. Briefly, fastq files from the same samples were combined and adaptors were trimmed using cutadapt (Martin, 2011). Parameters for read 2 trimming were 5’ adaptor sequence as ’GGCCAGTTTAAACTT’, 3’ adaptor sequence as ’GCATGGC’, maximum error rate of 0.2 and 27 base pair in length for trimmed read2. Trimmed paired reads were merged using FLASh with minimum overlap of 12 base pairs and maximum mismatch rate of 0.25 (Magoč and Salzberg, 2011). The resulting barcode is 27 base pairs including 12 bp barcode, 6 bp SphI restriction site and 9 bp UMIs. The barcode-UMI combinations with perfect match to all possible barcode- UMI combinations from the designed libraries were counted for analysis described below.

### Fitness calculation

Processed counts from each competition experiment of barcode-UMI combinations were combined with generation time estimated from optical density at each timepoint during fitness competition to calculate fold-change values using DESeq2 (Love et al., 2014). A minimum filter of 500 reads across 18 samples, including six time points in three replicates, was set for each barcode-UMI combination. The editing effect of each barcode-UMI combination was modeled as described previously by estimating the effect of generation time on the log fraction of barcode-UMI counts, with the Deseq2 design formula as follows (Sharon et al., 2018):

Counts ∼ Generation + Flask

Where “Counts” represent read counts of each barcode-UMI combination; “Generation” represents the number of generations from the start of the growth competition, estimated by optical density as described above; “Flask” indicate the flask replicate from which the sample originated. Log2 fold-change was estimated for counts per UMI across generation time for each barcode-UMI combination by Deseq2 (Love et al., 2014).

### Outlier removal and GxE fitness modeling

Individual UMI log2 fold changes (logFC) for the same variants were combined to estimate the variant fitness effect through a weighted least squares model using a custom Python script (modified from Ang *et al*, in submission). For each genomic editing guide-donor pair with associated barcode, we removed outlier barcode-UMIs with large median absolute deviations (MAD) from the median logFC for that barcode (logFC >3.5 x MAD from median logFC for that barcode), as each barcode-UMI should reflect the same fitness effect of the genomic edit. This was intended to remove barcode-UMIs which were in strains which had acquired off-target mutations during transformation, editing or growth, particularly strong beneficial de novo mutations which would otherwise skew fitness measurements. Next, for the ergosterol library we calculated the standard deviation of the logFC of the barcode-UMIs for each programmed barcode and removed programmed barcodes with logFC standard deviation greater than or equal .05, to remove highly variable barcodes not accounted for in the previous outlier detection step. We omitted this step for the QTL pools due to higher variance that was expected among very high effect variants in those pools. We then calculated inverse-variance weights for each barcode-UMI based on its read depth across the competition by fitting a regression model fitness standard deviation ∼ baseMean to the neutral UMIs at different read depths. This led to barcode-UMIs with more reads being weighted more highly, reflecting the higher confidence of their fitness effect measurements. Using Deseq2 variance directly gave inaccurately low estimates of variance. Genetic variant fitness effects across all competitions were then fit into a weighted least squares model using the weights mentioned above. The dependent variable was barcode-UMI logFC as measured by Deseq2, and the independent variables where the variant, the condition for the competition, and the interaction term between the variant and condition:

logFC ∼ variant + condition + variant:condition

The variant and condition terms were categorical variables (as was the variant:condition term), reflecting the variant the UMIs are linked to, and the growth competition the logFC values came from. The variant fitness in any given condition thus reflects the difference between the neutral barcode-UMIs in that condition and the variant-linked barcode-UMIs, weighted by the read depth of each UMI. Significance is determined by a weighted t-test, and p-values were adjusted for multiple testing using the Benjamini-Hochberg procedure and significant fitness effects were controlled at FDR = 0.01 (Benjamini and Hochberg, 1995).

### Fluconazole Ecological Enrichment Test

To test whether strains from particular ecological origins were enriched for variants with significant effects in a particular direction in fluconazole, we first split the variants with significant fitness effects in fluconazole into positive and negative effect variants. We then checked for each strain in the 1,011 yeast genomes if they were homozygous or heterozygous for the alternate allele we edited in at each significant variant. For positive effect variants, strains with the alternate allele had 1 added to their score, and for negative effect alleles, strains with the alternate allele had 1 subtracted from their score. The total number of negative effect variants was added to this score for all strains, as any strain with the reference allele for those sites in effect had the positive effect allele. The 1,011 yeast strains were then sorted by this score, and the top 100 were chosen to look at their ecological origins, as they were presumably the strains with the most evidence for being under selection for increased growth in fluconazole. A hypergeometric test was performed to determine enrichment of the top categories, “Human” and “Human, clinical.”

### Detecting significant GxE interactions

The weighted least squares model described above detected variants with significant fitness effects in a given condition as well as variants with gene-by-environment effects; all pairwise differences in fitness effects between conditions (e.g. 15 differences for variants measured in six conditions) were calculated. The p-values were adjusted for multiple testing using the Benjamini-Hochberg procedure and significant differences were controlled at FDR = 0.01 (Benjamini and Hochberg, 1995). At this threshold, none of the neutral/non-cutting variants exhibited GxE interactions.

### Permutation test for nonrandom clustering of GxE promoter variants

In order to test whether the level of clustering observed for hits in each annotation category at each distance was more than would be expected by chance, we permuted the hits 1000 times by choosing random variants of the same annotation to be hits, choosing the same number of promoter hits as exist in the real dataset, and then performing the same cluster analysis for these permuted sets. We then counted how many of these permuted data sets had greater than or equal to the number of genomic loci in clusters as the real data set to determine permutation p-values. For the final p- value for promoter variants at 8 bp, we performed 5000 permutations twice (two seeds). We then took the average of the two permutation p-values.

### Clonal genotyping

Single CRISPEY-BAR oligonucleotides containing partial sequence containing 5’ homology to pSAC200, 12 bp programmed barcode, restriction site region for cloning, 108 bp donor template sequence and 34 bp constant region were ordered from IDT as dsDNA eblocks for individual validation of genotype correct strains. The eblocks were amplified using primer #576 as forward primer and donor specific primers that append the 20 bp guide sequence and 3’ homology to pSAC200 to the eblocks. The resulting PCR products were bead purified and cloned into pSAC200, ligated with UMI-containing insert and transformed into yeast as described for library cloning above. The yeast transformants were induced for editing by culturing in 5 mL YNB -HIS -URA 2% raffinose media for 24 hr, passaged twice in 5 mL YNB -HIS -URA 2% galactose media for 24 hr each, then streaked out on YNB -URA 2% glucose (1.7g/L yeast nitrogen base (RPI); 5 g/L Ammonium Sulfate (ACROS organics); 1.9 g Dropout synthetic mix minus uracil, w/o nitrogen base (US Biological) and 20 g/L glucose (Sigma) 2% agar plates to obtain single edited clones. plasmids were cured from edited clones by restreaking on YNB 2% glucose 2% agar plates with 1 g/L 5-Fluororotic acid monohydrate (GoldBio). The single plasmid-cured colonies were amplified by growing in YNB 2% glucose media overnight and stored in YNB 2% glucose media with 15% glycerol at -80°C.

Colonies were streaked out from the frozen stock and lysed with Zymolyase 20T (US Biological) solution in 50 mM potassium phosphate buffer, pH 7.5. Cell lysates were used for genotyping using EmeraldAmp MAX PCR Mastermix (Takara), with primers #261 and #262 for determining barcode-UMI sequence and locus-specific primers (**Supplemental Table S1**). PCR cycles had an initial denaturation of 95°C for 2 min; then 35 cycles of 95°C for 10 s, 60°C for 15 s, 72°C for 20 s; then a final extension of 72°C for 5 min. PCR products were purified, Sanger sequenced and aligned to the reference genome using SGD BLAST to confirm the intended genotype (Cherry et al., 2012; Engel et al., 2014). For the QTL pools, colonies were randomly picked from edited cells plated on non-selective media after plasmid removal. Genomic amplicons of loci containing the associated variant edit were Sanger sequenced from barcoded colonies to calculate the editing rates shown in **Figure 1d**.

### qRT-PCR

Strains containing the Sanger sequencing-verified genotypes were thawed from frozen stock and grown overnight in 5 mL YNB 2% glucose media. 0.5 mL of the overnight culture was passaged to 50 mL YNB 2% glucose media with or without 30 mg/mL lovastatin. Cells were harvested after 5 generations of growth in media, approximately 12 hr after passaging. RNA was extracted by vortexing with 500 uL glass beads and 1 mL Trizol (Invitrogen) by manufacturer’s instructions. 8 ug total RNA of each sample were digested with RQ1 DNase for 1 hr at 37°C as manufacturer’s instructions and purified by overnight ethanol precipitation. 400 ng of the purified RNA from each sample were converted to cDNA using Superscript IV First-Strand Synthesis system by manufacturer’s instructions. qPCR was performed as described previously (Sharon et al., 2018). qPCR primers for ERG4 and ACT1 are included in Supplementary Table S1.

### Fitness validation

Strains containing the Sanger sequencing verified genotypes were thawed from frozen stock and grown overnight in SC media and mixed with the GFP control strain in 1 mL SC media with specified conditions in a 96-well plate. Cells were passaged every 12 hr and diluted to fresh 1 mL SC media with specified conditions. Six timepoints (T1-T6) were harvested once passage was complete. Harvested cells were spun down and resuspended in 1x DPBS (Gibco) and stored at 4°C and assayed by flow cytometry within 12 hr post-harvest. Generation time was estimated by measuring OD600 of the culture containing ZRS111 and GFP control strain at every time point. Competition for each edited strain against GFP control strain was replicated four times in four different wells, to control for spontaneous mutation during competition.

Ratios between each edited strain against GFP control strain were determined by flow cytometry assay, using an Attune NxT Flow Cytometer and Autosampler (ThermoFisher Scientific). GFP was detected using a 530 nm band-pass filter (BL1) with a 488 nm laser. The channel voltages were adapted from a previous study and set as follows: FSC: 200; SSC: 320; and BL1:480^41^. A threshold for FSC of 2.5 x 10^3^ A.U. was applied to exclude non-yeast events. Data analysis was performed using Attune NxT Software v2.7.

Doublets were removed by FSC gating and cell counts for GFP control strain were determined by BL1 gating and the remaining cells were counted as the non-fluorescent, corresponding to edited strains. Samples with fewer than 500 total cells gated, as well as samples with cell counts of less than 3 for either GFP or edited strains, were excluded.

Log2 ratios between edited strain count and GFP control strain count were calculated for each sample and fitted to a slope for the estimated generations within each replicate.

The slopes were normalized by subtracting the slope calculated by the competition of a non-variant edit, barcode-only control to the GFP control strain in the same replicate.

Finally, the mean and standard error for slopes across four replicates were calculated for each edited strain, representing pairwise fitness values.

### Data and Software Availability

All raw sequencing data have been deposited in the NCBI Sequence Read Archive ”PRJNA827354”. All software and code used for the design and barcode-UMI count analysis are available upon request. Code used to create specific figures is available upon request. Sequences for oligonucleotides used in this study can be found in Table S1. Sanger sequencing data related to Figure 1e can be found at: Chen, Shi-An (2022), “Gene-by-environment interactions are pervasive among natural genetic variants”, Mendeley Data, V1, doi: 10.17632/sm32n7ms8h.1

## Supplementary Tables

**Table S1. Plasmid sequences, primers and oligonucleotides used in this study.** This table contains 1) plasmid sequence used in this study; 2) primers used in this study; gblock oligonucleotides for individual validation used in this study; 4-7) synthesized oligonucleotide pools for CRISPEY-BAR used in this study, including pools for cobalt chloride, caffeine, fluconazole and ergosterol pathway.

**Table S2. Assignment of array synthesized oligonucleotides for each CRISPEY- BAR pool by well number.** This table contains information for well pooling information, with each column contains the well IDs from synthesized oligonucleotide array assigned to each well.

